# *PHAROH* lncRNA regulates c-Myc translation in hepatocellular carcinoma via sequestering TIAR

**DOI:** 10.1101/2021.03.23.436634

**Authors:** Allen T. Yu, Carmen Berasain, Sonam Bhatia, Keith Rivera, Bodu Liu, Frank Rigo, Darryl J. Pappin, David L. Spector

## Abstract

Hepatocellular carcinoma, the most common type of liver malignancy, is one of the most lethal forms of cancer. We identified a long non-coding RNA, *Gm19705*, that is over-expressed in hepatocellular carcinoma and mouse embryonic stem cells. We named this RNA *<u>P</u>luripotency and <u>H</u>epatocyte <u>A</u>ssociated <u>R</u>NA <u>O</u>verexpressed in <u>H</u>CC*, or *PHAROH*. Depletion of *PHAROH* impacts cell proliferation and migration, which can be rescued by ectopic expression of *PHAROH*. RNA-seq analysis of *PHAROH* knockouts revealed that a large number of genes with decreased expression contain a *c-Myc* motif in their promoter. C-MYC is decreased at the protein level, but not the mRNA level. RNA-antisense pulldown identified nucleolysin TIAR, a translational repressor, to bind to a 71-nt hairpin within *PHAROH*, sequestration of which increases c-MYC translation. In summary, our data suggest that *PHAROH* regulates c-MYC translation by sequestering TIAR and as such represents a potentially exciting diagnostic or therapeutic target in hepatocellular carcinoma.

## Introduction

Hepatocellular carcinoma (HCC), the most common type of liver malignancy, is one of the most lethal forms of cancer (Asrani et al., 2019). HCC is the fifth-most frequently diagnosed cancer and the third-leading cause of cancer-related deaths worldwide (Villanueva, 2019). The molecular landscape of HCC is very complex and includes multiple genetic and epigenetic modifications which could represent new diagnosis and therapeutic targets. In this sense, multiple studies have established molecular classifications of HCC subtypes that could be related to clinical management and outcomes (Dhanasekaran et al., 2019; Llovet et al., 2018). For instance, Hoshida et al. classified HCC into S1, S2, and S3 subtypes by means of their histological, pathological, and molecular signatures (Hoshida et al., 2009). S1 tumors exhibit high TGF-β and Wnt signaling activity but do not harbor mutations or genomic changes. The tumors are relatively large, poorly-differentiated, and associated with poor survival. S2 tumors have increased levels of c-Myc and phospho-Akt and overexpress α-fetoprotein, an HCC serum biomarker. S3 tumors harbor mutations in *CTNNB1* (β-catenin) but tend to be well-differentiated and are associated with good overall survival.

The standard of care for advanced HCC is treatment with sorafenib, a multi-kinase inhibitor that targets Raf, receptor tyrosine kinases (RTKs), and the platelet-derived growth factor receptor (PDGFR). Sorafenib extends the median survival time from 7.9 months to 10.7 months, and lenvatinib, a multiple VEGFR kinase inhibitor, has been reported to extend survival to 13.6 months (Llovet et al., 2018; Philip et al., 2005; Rimassa & Santoro, 2009). Combination therapies of VEGF antagonists together with sorafenib or erlontinib are currently being tested (Dhanasekaran et al., 2019; Greten et al., 2019; Quintela-Fandino et al., 2010). However, even with the most advanced forms of treatment, the global death toll per year reaches 700,000, creating a mortality ratio of 1.07 with a 5-year survival rate of 18% (Ferlay et al., 2010; Siegel et al., 2014; Villanueva, 2019). Not only is it difficult to diagnose HCC in the early stages, but there is also a poor response to the currently available treatments. Thus, novel therapeutic targets and treatments for HCC are urgently needed.

The ENCODE consortium revealed that as much as 80% of the human genome can be transcribed, while only 2% of the genome encodes for proteins (Djebali et al., 2012). Thousands of transcripts from 200 nucleotides (nt) to over one-hundred kilobases (kb) in length, called long non-coding RNAs (lncRNAs), are the largest and most diverse class of non-protein-coding transcripts. They commonly originate from intergenic regions or introns and can be transcribed in the sense or anti-sense direction. Most are produced by RNA polymerase II and can be capped, spliced and poly-adenylated (reviewed in Rinn & Chang, 2012). Strikingly, many are expressed in a cell or tissue-specific manner and undergo changes in expression level during cellular differentiation and in cancers (Costa, 2005; Dinger et al., 2008). These lncRNAs present as an exciting class of regulatory molecules to pursue, as some are dysregulated in HCC and have potential to be specific to a subtype of HCC (Li et al., 2015).

One of the few examples of a lncRNA that has been studied in the context of HCC is the homeobox (HOX) anti-sense intergenic RNA (*HOTAIR*). This transcript acts in trans by recruiting the Polycomb repressive complex 2 (PRC2), the lysine-specific histone demethylase (LSD1) and the CoREST/REST H3K4 demethylase complex to their target genes (Ezponda & Licht, 2014). *HOTAIR* promotes HCC cell migration and invasion by repressing RNA binding motif protein 38 (RBM38), which is otherwise targeted by p53 to induce cell cycle arrest in G1 (Shu et al., 2006; Yu et al., 2015). Another mechanism through which lncRNAs function involves inhibitory sequestration of miRNAs and transcription factors (Cesana et al., 2011). In HCC, the lncRNA *HULC* (highly upregulated in liver cancer) sequesters *miR-372*, which represses the protein kinase PRKACB, and down-regulates the tumor suppressor gene *CDKN2C* (p18) (Wang et al., 2010). Similarly, the highly-conserved *MALAT1* lncRNA controls expression of a set of genes associated with cell proliferation and migration and is upregulated in many solid carcinomas (Amodio et al., 2018; Lin et al., 2007); siRNA knockdown of *MALAT1* in HCC cell lines decreases cell proliferation, migration, and invasion (Lai et al., 2012).

Only a small number of the thousands of lncRNAs have been characterized in regard to HCC. Therefore, whether and how additional lncRNAs contribute to HCC remains unknown, and it is not fully understood how lncRNAs acquire specificity in their mode of action at individual gene loci. A lack of targetable molecules limits the effectiveness of treatments for HCC, and this class of regulatory RNAs has great potential to provide novel therapeutic targets.

Here, we reanalyzed naïve and differentiated transcriptomes of mouse embryonic stem cells (ESCs) in the context of the GENCODE M20 annotation. We aimed to identify lncRNAs that are required for the pluripotency gene expression program, and dysregulated in cancer, with a specific focus on HCC. Since normal development and differentiation are tightly regulated, dysfunction of potential regulatory RNAs may lead to various disease phenotypes including cancer. One lncRNA that is highly upregulated in HCC is of special interest, and we show that it interacts with and sequesters the translation repressor nucleolysin TIAR resulting in an increase of c-Myc translation. Together, our findings identified a mechanism by which a lncRNA regulates translation of c-MYC in HCC by sequestering a translation inhibitor and as such has potential as a therapeutic target in HCC.

## Results

### Deep sequencing identifies 40 long non-coding RNAs dysregulated in embryonic stem cells and cancer

Since normal development and differentiation are tightly regulated processes, we reasoned that lncRNAs whose expressions are ESC specific and can be found to also exhibit altered expression in cancer, may have important potential roles in regulating critical cellular processes.

We re-analyzed the raw data from our published differential RNA-seq screen comparing lncRNA expression in mouse ESCs vs neural progenitor cells (NPCs) (Bergmann et al., 2015), using updated bioinformatic tools and the recently released GENCODE M20 annotation (January 2019), which has nearly 2.5 times more annotated lncRNAs than the previously used GENCODE M3. Principal component analysis (PCA) of the processed data showed that ESCs and NPCs independently cluster, and the difference between ESC cell lines (AB2.2) and mouse derived ESCs only accounted for 4% of the variance (Fig. 1A). Additionally, we prioritized transcripts with an FPKM value greater than 1, and those that were more than 2-fold upregulated in ESCs compared to NPCs. This left us with 147 ESC specific transcripts. Since our goal is to discover novel transcripts that may play a role in the progression of human cancer, we first needed to identify the human homologues of the 147 mouse ESC transcripts. In addition to sequence conservation, we also evaluated syntenic conservation of the mouse lncRNAs to the human genome, due to the fact that many lncRNAs are not conserved on the sequence level. Finally, we queried TCGA databases via cBioportal, to find lncRNAs that were altered in cancer (Fig. 1B). A final candidate list of 40 lncRNAs that are enriched in ESCs, and dysregulated in cancer, was identified (Table 1). Our candidate list contains lncRNAs that have a wide range of expression, and also contains several previously identified lncRNAs that have been found to be dysregulated in cancer (*NEAT1*, *FIRRE*, *XIST*, *DANCR*, and *GAS5*), verifying the validity of the approach (Fig. S1A) (Ji et al., 2019; Soudyab et al., 2016; Yuan et al., 2016).

**Figure 1.**
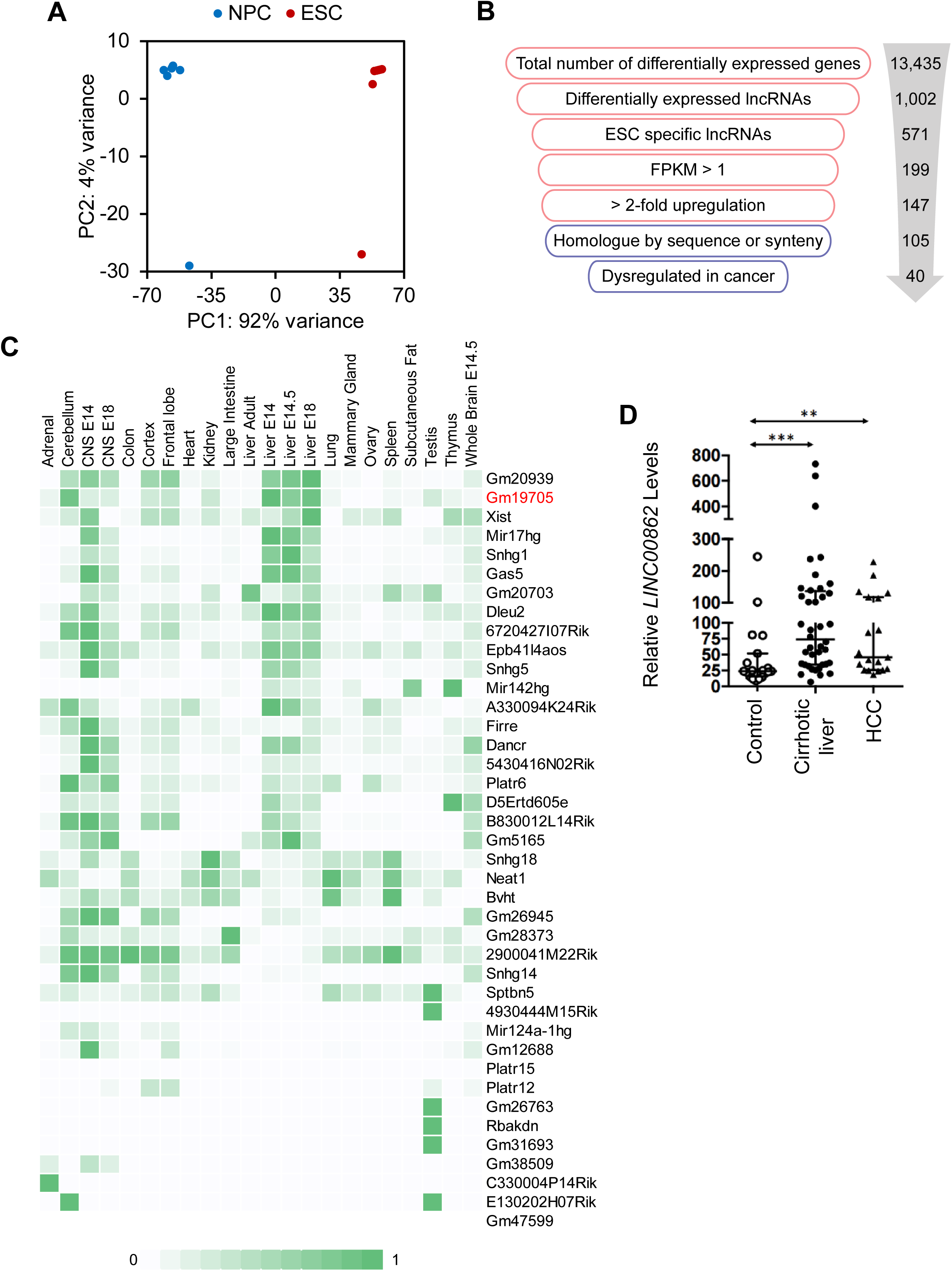
LncRNA screen to identify transcripts enriched in ESCs and dysregulated in cancer. A. PCA plot of 10 RNA-seq libraries from mouse derived ESCs, and two from cell lines. Differentiation from ESCs to NPCs created the largest difference in variance, while there was minimal difference between isolated clones vs. cell lines. B. Workflow of the filtering process performed to obtain ESC enriched lncRNAs that are also dysregulated in cancer. Red indicates analysis performed in mouse and blue indicates human. C. LncRNA candidate expression across ENCODE tissue datasets show that lncRNAs are mostly not pan-expressed, but are rather tissue specific. Counts are scaled per row. D. *LINC00862* is upregulated in both human cirrhotic liver and HCC tumor samples when compared to control patient liver tissue samples. **p < 0.01; ***p < 0.005; Student’s t-test.

**Table 1.**
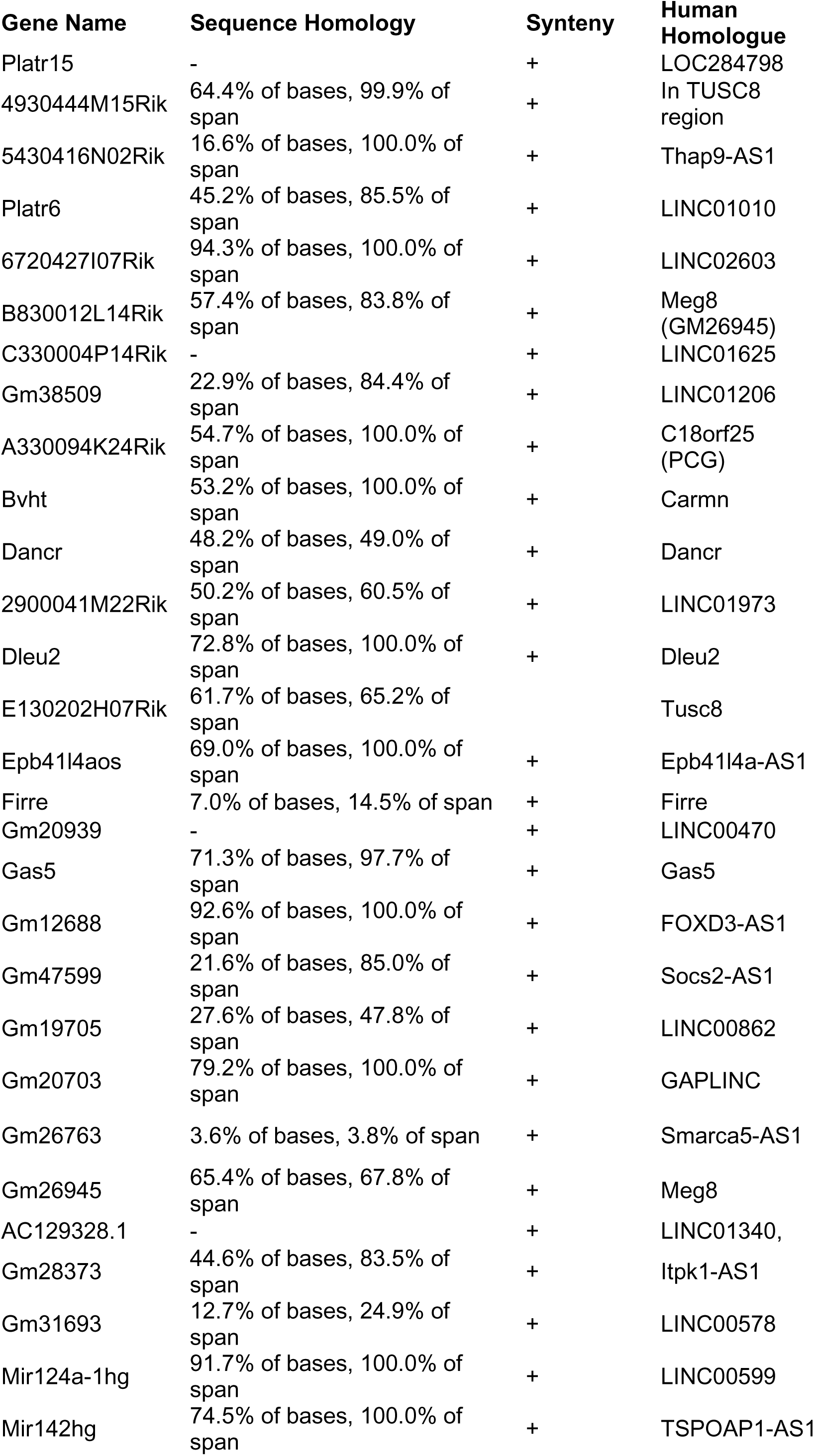
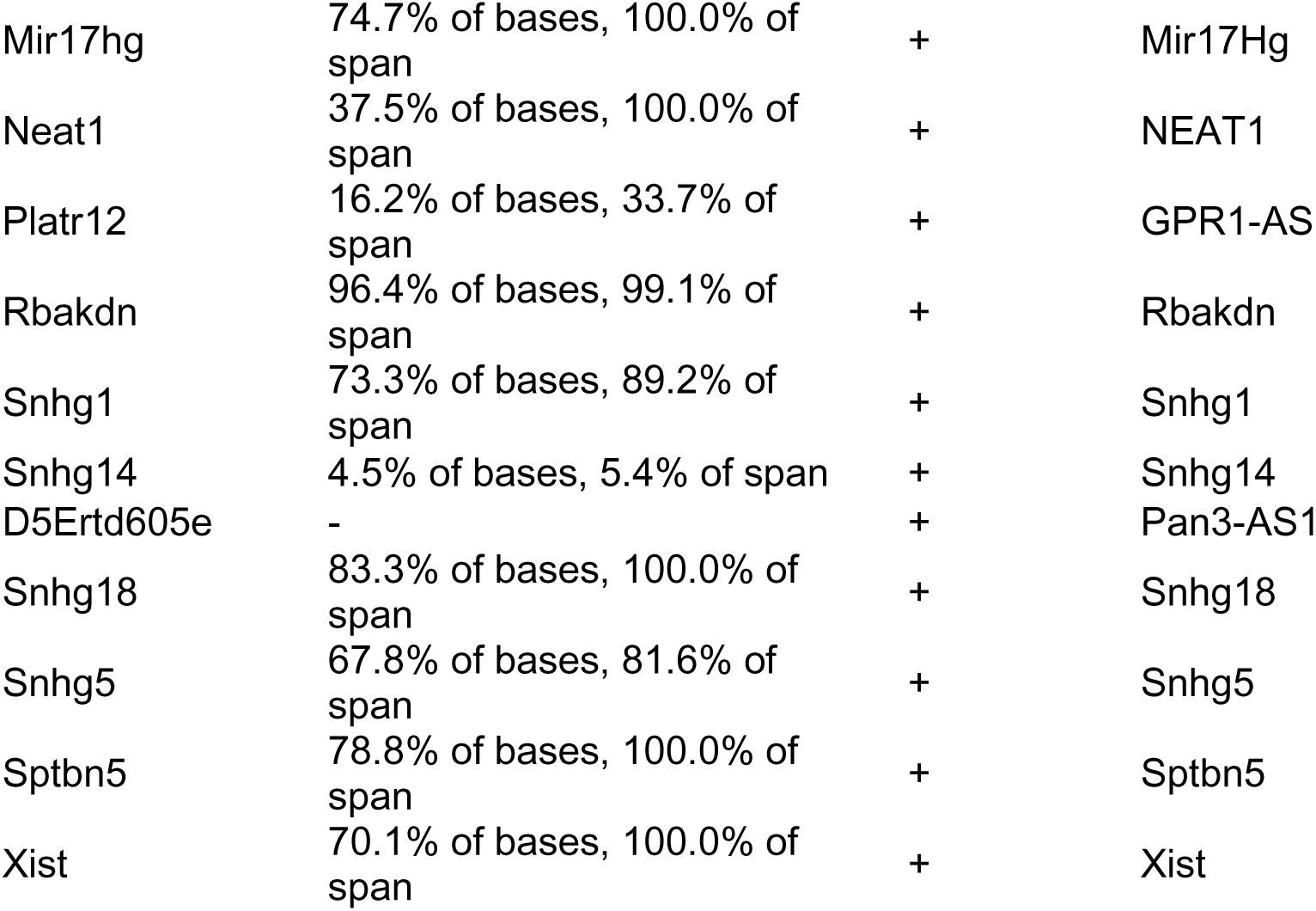
Candidate list of lncRNAs that are enriched in ESCs and dysregulated in cancer.

| Gene Name | Sequence Homology | Synteny | Human Homologue |
| --- | --- | --- | --- |
| Platr15 | - | + | LOC284798 |
| 4930444M15Rik | 64.4% of bases, 99.9% of span | + | In TUSC8 region |
| 5430416N02Rik | 16.6% of bases, 100.0% of span | + | Thap9-AS1 |
| Platr6 | 45.2% of bases, 85.5% of span | + | LINC01010 |
| 6720427I07Rik | 94.3% of bases, 100.0% of span | + | LINC02603 |
| B830012L14Rik | 57.4% of bases, 83.8% of span | + | Meg8 (GM26945) |
| C330004P14Rik | - | + | LINC01625 |
| Gm38509 | 22.9% of bases, 84.4% of span | + | LINC01206 |
| A330094K24Rik | 54.7% of bases, 100.0% of span | + | C18orf25 (PCG) |
| Bvht | 53.2% of bases, 100.0% of span | + | Carmn |
| Dancr | 48.2% of bases, 49.0% of span | + | Dancr |
| 2900041M22Rik | 50.2% of bases, 60.5% of span | + | LINC01973 |
| Dleu2 | 72.8% of bases, 100.0% of span | + | Dleu2 |
| E130202H07Rik | 61.7% of bases, 65.2% of span |  | Tusc8 |
| Epb41l4aos | 69.0% of bases, 100.0% of span | + | Epb41l4a-AS1 |
| Firre | 7.0% of bases, 14.5% of span | + | Firre |
| Gm20939 | - | + | LINC00470 |
| Gas5 | 71.3% of bases, 97.7% of span | + | Gas5 |
| Gm12688 | 92.6% of bases, 100.0% of span | + | FOXD3-AS1 |
| Gm47599 | 21.6% of bases, 85.0% of span | + | Socs2-AS1 |
| Gm19705 | 27.6% of bases, 47.8% of span | + | LINC00862 |
| Gm20703 | 79.2% of bases, 100.0% of span | + | GAPLINC |
| Gm26763 | 3.6% of bases, 3.8% of span | + | Smarca5-AS1 |
| Gm26945 | 65.4% of bases, 67.8% of span | + | Meg8 |
| AC129328.1 | - | + | LINC01340, |
| Gm28373 | 44.6% of bases, 83.5% of span | + | Itpk1-AS1 |
| Gm31693 | 12.7% of bases, 24.9% of span | + | LINC00578 |
| Mir124a-1hg | 91.7% of bases, 100.0% of span | + | LINC00599 |
| Mir142hg | 74.5% of bases, 100.0% of span | + | TSPOAP1-AS1 |
| Mir17hg | 74.7% of bases, 100.0% of span | + | Mir17Hg |
| Neat1 | 37.5% of bases, 100.0% of span | + | NEAT1 |
| Platr12 | 16.2% of bases, 33.7% of span | + | GPR1-AS |
| Rbakdn | 96.4% of bases, 99.1% of span | + | Rbakdn |
| Snhg1 | 73.3% of bases, 89.2% of span | + | Snhg1 |
| Snhg14 | 4.5% of bases, 5.4% of span | + | Snhg14 |
| D5Ertd605e | - | + | Pan3-AS1 |
| Snhg18 | 83.3% of bases, 100.0% of span | + | Snhg18 |
| Snhg5 | 67.8% of bases, 81.6% of span | + | Snhg5 |
| Sptbn5 | 78.8% of bases, 100.0% of span | + | Sptbn5 |
| Xist | 70.1% of bases, 100.0% of span | + | Xist |

We analyzed the ENCODE expression datasets of adult mouse tissue to compare the expression levels of the candidates across tissues (Fig. 1C). LncRNAs are known to have distinct expression patterns across different tissues, and our results support the notion that lncRNAs are generally not pan-expressed. Interestingly, many of the identified lncRNAs are enriched in embryonic liver, which is the organ with the most regenerative capacity, yet never grows past its original size.

From here, we decided to focus on liver enriched candidate mouse lncRNAs, especially those that were primarily dysregulated in liver cancers. Because HCC is one of the deadliest cancers and has inadequate treatment options, we focused on lncRNAs that were dysregulated in HCC, *LINC00862*, *TSPOAP-AS1*, *MIR17HG*, and *SNHG5*, with their mouse counterparts being *Gm19705*, *Mir142hg*, *Mir17hg*, and *Snhg5*, respectively. Out of these four lncRNAs that were detected to be amplified in HCC, *LINC00862* was the highest at 13% of all liver cancer cases (Fig. S1B). We assayed LINC00862 expression in human samples obtained from healthy and cirrhotic livers and HCC nodules. Indeed, we found that levels of *LINC00862* were elevated in HCC tumor nodules, but also in cirrhotic liver, suggesting that it may play a role in HCC progression (Fig. 1D).

In order to use a more tractable model system, we assessed the conservation of *LINC00862* and its potential mouse counterpart, *GM19705*, which was internally designated as *lnc05* in previous analyses (Bergmann et al., 2015). While much shorter, *GM19705* has 51% sequence identity and the gene order is syntenically conserved, although a reversal event most likely occurred within the locus (Fig S1C). Weighted gene correlation network analysis of *GM19705* identified that its expression is highly correlated with those of cell cycle genes, such as *BRCA1* and *BRCA2* (Fig. S1D). GO-term analysis of the cluster identified cell cycle processes as highly enriched, indicating that *GM19705* may play a role in the regulation of the cell cycle (Fig. S1E). Re-analysis of previously published single cell analysis of normal adult mouse liver (Tabula Muris et al., 2018) identified *GM19705* expression to be low overall, as expected, but highly expressed exclusively in a subset of hepatocytes (Fig. S1F).

Our analysis identified *GM19705*/*LINC00862* as a lncRNA that is expressed in ESCs and dysregulated in HCC. We found that *GM19705* is also highly expressed in developing liver and exclusively in adult hepatocytes, and it may have a potential function to regulate the cell cycle. Therefore, we named this mouse lncRNA – *<u>P</u>luripotency and <u>H</u>epatocyte <u>A</u>ssociated <u>R</u>NA <u>O</u>verexpressed in <u>H</u>CC*, or *PHAROH*.

### PHAROH is a novel lncRNA that is highly expressed in embryonic liver and mouse hepatocellular carcinoma

*PHAROH* is an intergenic lncRNA located on mouse chr1:1qE4. 5’ and 3’ rapid extension of cDNA ends (RACE) revealed the presence of two isoforms that share two common exons and are both ∼450 nt (Fig. 2A). In silico analysis of the coding potential by three independent algorithms, which use codon bias (CPAT/CPC) and comparative genomics (PhyloCSF), all point towards the low coding potential score of *PHAROH*, compared to the *Gapdh* control (Fig. S2A-B). From here on, only qPCR primers that amplify common exons were used. We confirmed expression levels of *PHAROH* in developing liver by assaying the liver bud from E14 and E18 embryos and found that they were 7-9 fold enriched compared to adult liver (Fig. 2B). Because the liver is one of the main sites of hematopoiesis in the embryo, we measured *PHAROH* levels in embryonic blood and found that expression was exclusive to the liver, and not to hematopoietic cells (Fig. S2C). *PHAROH* was also found to be upregulated in a partial hepatectomy model of liver regeneration (Fig. S2D), where expression was correlated with time points of concerted cell division, but did not fluctuate across the cell cycle (Fig. S2E). To confirm *PHAROH*’s involvement in HCC, we used a diethylnitrosamine (DEN) induced carcinogenic model of liver injury. By 11 months post DEN treatment, we were able to visualize HCC tumor nodules, which had elevated levels of *PHAROH* (Fig. 2C). In order to facilitate the molecular and biochemical study of *PHAROH*, we chose two mouse HCC cell lines, Hepa1-6 and Hepa1c1c7, and indeed found that *PHAROH* was 3-4-fold more enriched than in ESCs, and 8-10-fold increased over the AML12 mouse normal hepatocyte cell line (Fig. 2D).

**Figure 2.**
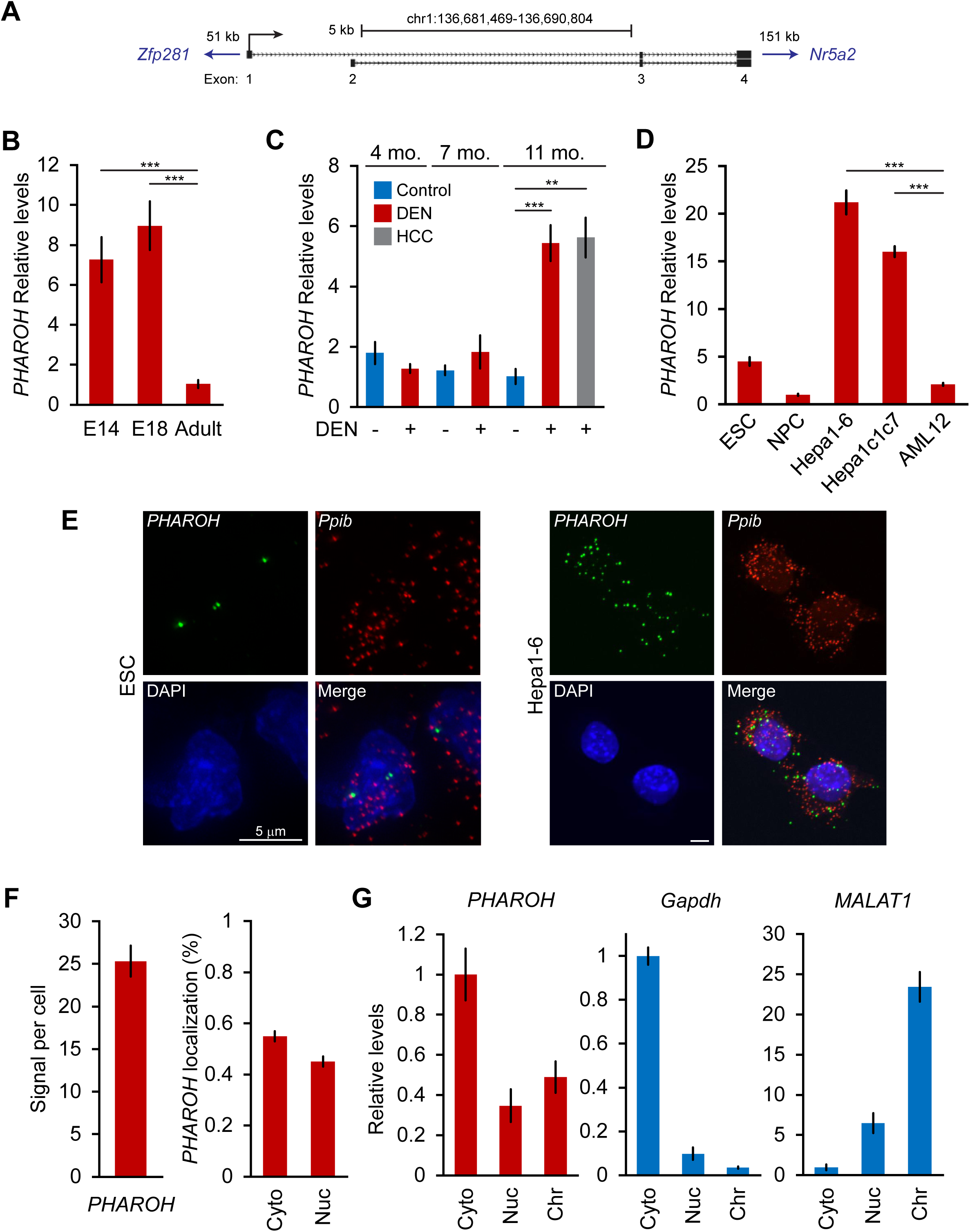
*PHAROH* lncRNA is highly expressed in ESCs, embryonic liver, models of hepatocarcinogenesis, and HCC cell lines. A. 5’ 3’ RACE reveals two isoforms for *PHAROH*, which have exons 3 and 4 in common. *PHAROH* is an intergenic lncRNA where the nearest upstream gene is *Zfp218* (51 kb away), and downstream is *Nr5a2* (151 kb away). B. *PHAROH* is highly expressed in embryonic liver in E14 and E18 mice, but not adult liver C. A DEN model of hepatocarcinogenesis shows high upregulation of *PHAROH* in the liver and HCC tumor nodules (gray bar) in DEN treated mice. D. *PHAROH* is upregulated in HCC cell lines (Hepa1-6, and Hepa1c1c7) compared to normal mouse hepatocytes (AML12). E. Single molecule RNA-FISH of *PHAROH* in ESCs shows nuclear localization and an average of 3-5 foci per cell. In Hepa1-6 cells, *PHAROH* shows 25 foci per cell, distributed evenly between the nucleus and cytoplasm (n=75 cells for each sample). F. Quantitation of panel *PHAROH* foci in panel E in HepA1-6 cells G. Cellular fractionation of Hepa1-6 cells shows equal distribution of *PHAROH* in the cytoplasm and nucleus, where it also binds to chromatin. *Gapdh* is predominantly cytoplasmic, and *MALAT1* is bound to chromatin. **p < 0.01; ***p < 0.005; Student’s t-test.

Single molecule RNA-FISH revealed that *PHAROH* is entirely nuclear in ESCs, with an average of 3-5 foci per cell, whereas it is evenly distributed between the nucleus and cytoplasm in Hepa1-6 cells, with an average of 25 foci per cell (Fig. 2E-F). Isoform 1 is expressed mostly in ESCs while isoform 2 of *PHAROH* dominates HCC cell lines (Fig. S2F). Cellular fractionation of Hepa1-6 cells corroborates the RNA-FISH determined localization of *PHAROH* as well, which *GAPDH* and *MALAT1* localized correctly to previously determined cellular fractions (Fig. 2G). Additional lncRNAs tested, such as *XIST*, *FIRRE*, and *NEAT1*, also localized to their expected cellular fractions (Fig. S2G). It was also determined that *PHAROH* has a relatively longer half-life in the Hepa1-6 cell line (10.8 h), compared to *MALAT1* (8.0 h), and *XIST* (4.2 h) (Fig. S2H) (Tani et al., 2012; Yamada et al., 2015). Taken together, *PHAROH* is an embryonic stem cell and fetal liver specific lncRNA, that is upregulated in the context of hepatocellular carcinoma.

### Targeted knockout of PHAROH

To evaluate the functional role of *PHAROH*, we generated targeted knockouts using CRISPR/Cas9 technology. Two sgRNA guides were designed to delete a region ∼700 bp upstream of the TSS, and ∼100 bp downstream of the TSS. We chose to transiently express enhanced specificity Cas9 (eSpCas9-1.1) in order to increase specificity, decrease off-target double stranded breaks, and also to avoid stable integration of Cas9 endonuclease due to its transformative potential (Slaymaker et al., 2016). In addition to using two guides targeting *PHAROH*, we used an sgRNA targeting renilla luciferase as a non-targeting control. Each guide was cloned into a separate fluorescent protein vector (GFP or mCherry) to allow for subsequent selection. Cells were single cell sorted 48h after nucleofection to account for heterogeneity of deletions among a pooled cell population, which may give certain cells a growth advantage. 85% of the cells were GFP+/mCherry+, and we selected four clones for subsequent analysis (Fig. S3A). All selected clones had the correct homozygous deletion when assayed by genomic PCR (Fig. 3A). qRT-PCR indicated that *PHAROH* was knocked down 80-95% (Fig. 3B).

**Figure 3.**
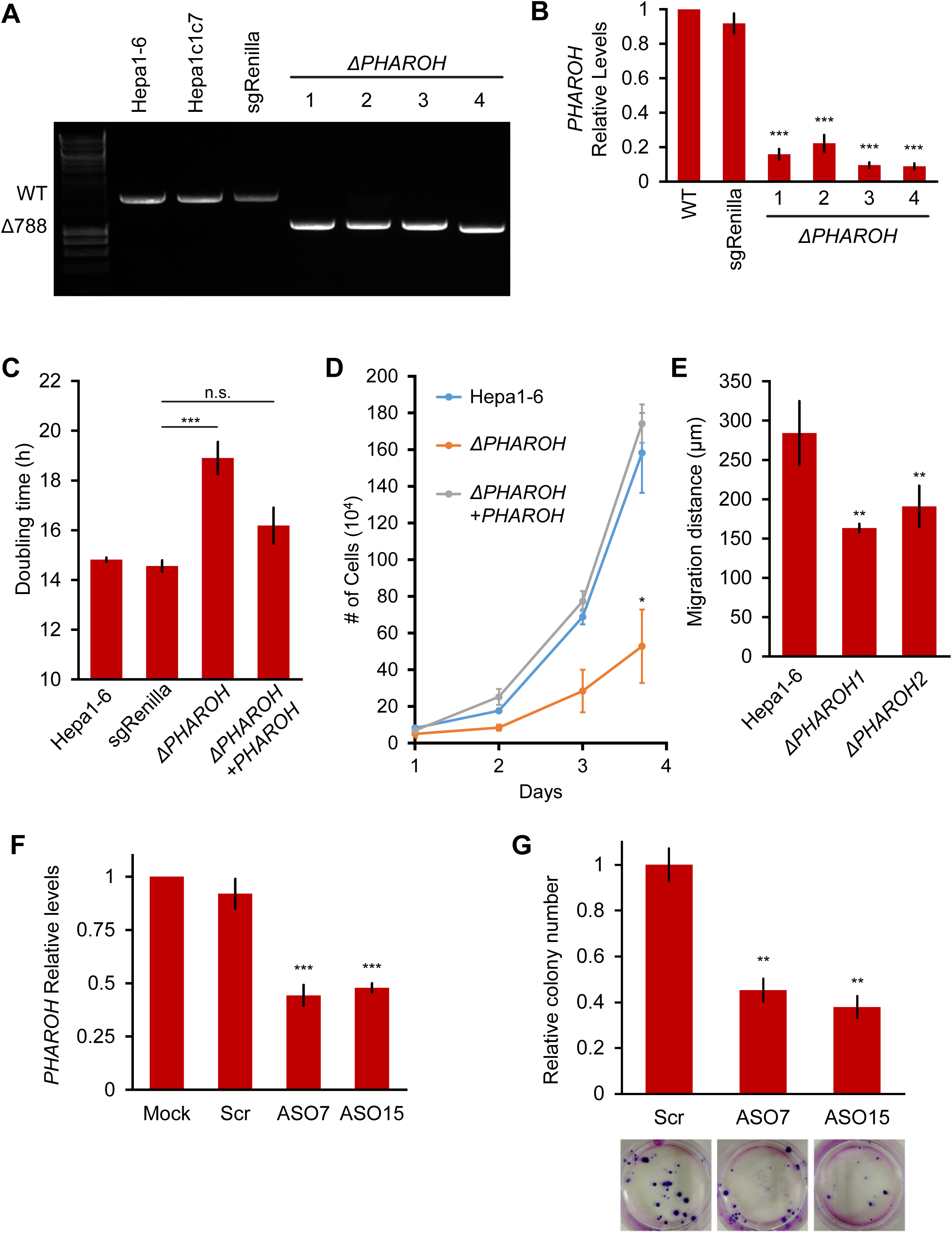
Depletion of *PHAROH* results in a proliferation defect. A. Four isolated clones all have a comparable deletion of 788 bp. The wildtype band is ∼1.8 kb. B. qRT-PCR of *PHAROH* knockout clones show a >80% reduction in *PHAROH* levels. C. Aggregated doubling time of clones shows knockout of *PHAROH* increases doubling time from 14.8h to 18.6h. Addition of *PHAROH* back into knockouts rescues this defect. D. Manual cell counting shows proliferation defect in *PHAROH* knockout cells that is rescued by ectopic expression of *PHAROH*. E. Migration distance for *PHAROH* knockout clones is decreased by 50%. F. 50% Knockdown of *PHAROH* can be achieved using both ASO7 and ASO15 at 24h. G. Colony formation assay of Hepa1-6 cells that are treated with scrambled or *PHAROH* targeting ASOs. After seeding 200 cells and two weeks of growth, a 50% reduction in relative colony number is observed. *p < 0.05; **p < 0.01; ***p < 0.005; Student’s t-test.

We assayed the proliferative state of the *PHAROH* knockout clones and found a decrease in proliferation. The doubling time of the knockout clones increased to 18.2h, compared to the wildtype doubling time of 14.8h, and ectopic expression of PHAROH reduced the doubling time to nearly widetype levels (Fig. 3C). Ectopic expression of *PHAROH* also successfully rescued the proliferation phenotype in the knockout clones, suggesting that *PHAROH* functions in *trans* (Fig. 3D). Migration distance was also decreased by 50% in the knockout clones (Fig. 3E).

In addition to assessing the role of *PHAROH* in knockout clones we also employed the use of antisense oligonucleotides (ASO) to knockdown *PHAROH*. We treated cells independently with a control scrambled cEt ASO, or two independent cEt ASOs complementary to the last exon of *PHAROH*. ASOs were nucleofected at a concentration of 2 uM, and we are able to achieve a >90% knockdown at 24h, and a ∼50% knockdown was still achieved after 96h (Fig. S3B). Proliferation assays using manual cell counts and MTS assay shows a 50% reduction in proliferation at 4 days (96h), similar to that achieved in our knockout clones, further supporting a role of *PHAROH* in cell proliferation (Fig. S3C). Addition of the ASO into the medium allowed for the knockdown to persist for longer duration to study the impact on clonogenic ability (Fig. 3F). Colony formation assays demonstrated that knockdown of *PHAROH* significantly inhibits clonogenic growth of HCC cells in a dose dependent manner (Fig. 3G, S3D).

To investigate the global effect of *PHAROH* depletion, we performed poly(A)+ RNA-seq on control and knockout clones (Fig. S4A, S4B). We identified 810 differentially expressed genes, and GO term analysis revealed regulation of cell proliferation, locomotion, and cell motility as the highest enriched terms (Fig. 4A). To determine if these differentially expressed genes were predominantly controlled by common transcription factors, we performed de novo and known motif analysis. Interestingly, promoter motif analysis of differentially expressed genes revealed enrichment of the c-Myc motif in our dataset suggesting a subset of the genes were under the transcriptional control of c-Myc (Fig. S4C). This was intriguing because c-Myc is known to regulate cell proliferation, and is highly amplified in nearly half of hepatocellular carcinomas (Zheng et al., 2017). However, c-Myc expression changes were not detected in our RNA-seq analysis, nor was there any statistically significant change compared to sgRenilla controls when assayed by qRT-PCR (Fig. 4B). Strikingly, *c-MYC* protein levels were substantially decreased in all of the *PHAROH* knockout clones, as detected by western blot and immunofluorescence, suggesting that *PHAROH* regulates *c-Myc* post-transcriptionally (Fig. 4C, S4D). qRT-PCR of genes downstream of c-Myc that were identified through our analysis were also significantly downregulated in *PHAROH* knockout clones (Fig. 4D). Thus, we suggest that depletion of *PHAROH* decreases *c-MYC* protein levels, and ultimately cell proliferation.

**Figure 4.**
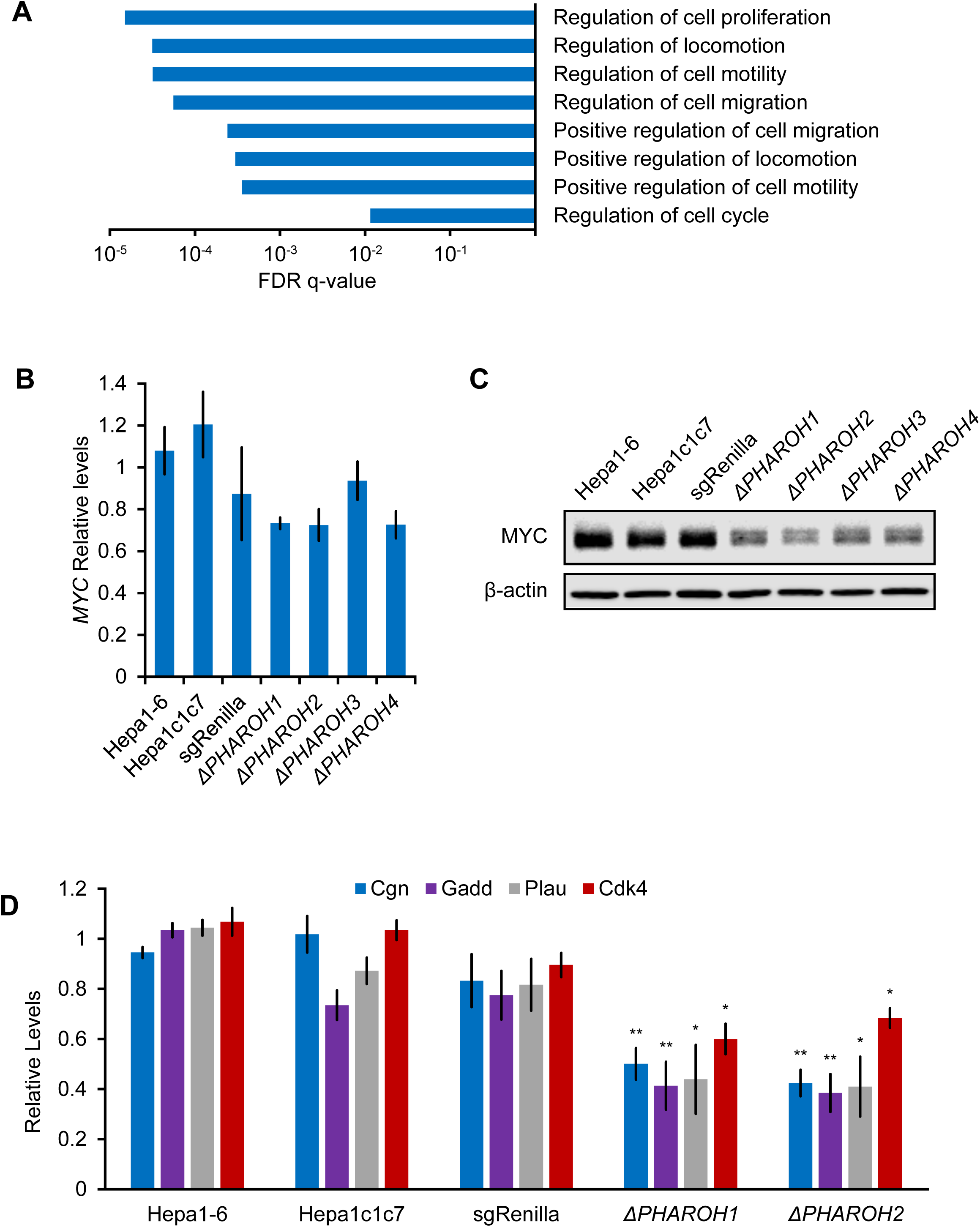
Gene expression analysis of *PHAROH* knockout cells reveals a link to c-MYC. A. GO term analysis of differentially expressed genes shows enrichment of cell proliferation and migration genes B. qRT-PCR of c-Myc mRNA levels indicate that c-Myc transcript does not appreciably change when *PHAROH* is knocked out. C. Western blot analysis of c-MYC protein shows downregulation of protein levels in *PHAROH* knockout cells. β-ACTIN is used as a loading control. D. qRT-PCR of genes downstream of c-Myc shows a statistically significant decrease in expression. *p < 0.05; **p < 0.01; Student’s t-test.

### RAP-MS identifies TIAR as the major interactor of PHAROH

LncRNAs can act as structural scaffolds to promote interaction between protein complexes or to sequester a specific protein (Lee et al., 2016; Tsai et al., 2010). Because modulation of *PHAROH* levels change *c-Myc* protein levels, but not mRNA levels to a significant degree, we hypothesized that *PHAROH* may be regulating the translation of *c-MYC* through a protein mediator. In order to search for *PHAROH* interacting proteins, we used a pulldown method adapted from the previously published RNA antisense purification-mass spectrometry (RAP-MS) (McHugh et al., 2015). In lieu of pooling all available antisense capture biotinylated oligonucleotides (oligos), we reasoned that individual oligos may be similarly effective, and can be used as powerful biological replicates. In addition, we would minimize oligo-specific off targets by verifying our results with multiple oligos. To this end, we screened through five 20-mer 3’ biotinylated DNA oligos that tiled the length of *PHAROH*, and found that four out of the five oligos pulled down >80% of endogenous *PHAROH*, while the pulldown of a control RNA, *PPIB*, remained low. (Fig. 5A, S5A).

**Figure 5.**
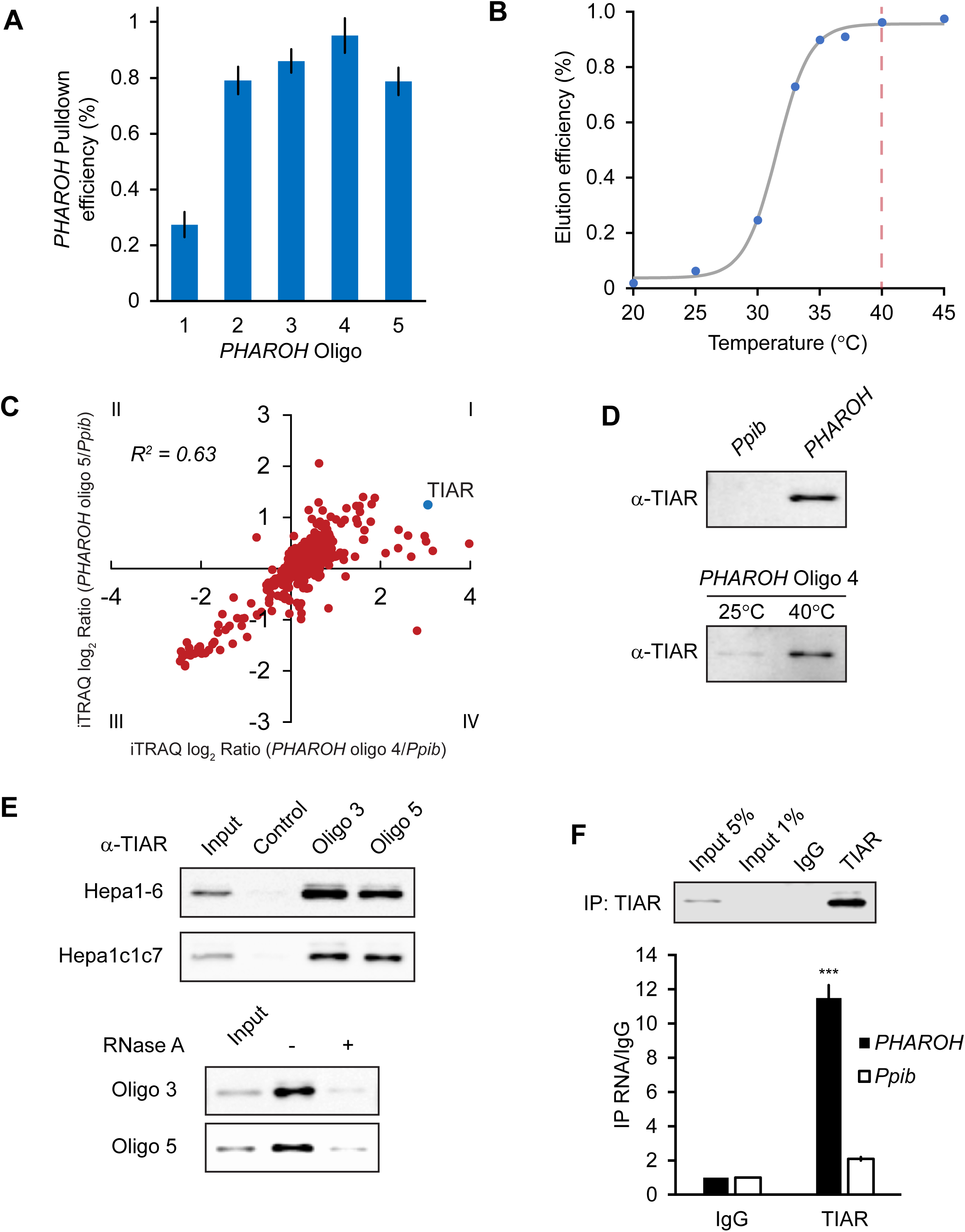
RAP-MS identifies TIAR as a major interactor of *PHAROH*. A. Five different biotinylated oligos antisense to *PHAROH* were screened for pulldown efficiency. Oligos 2-5 can pull down *PHAROH* at ∼80% efficiency or greater B. *PHAROH* can be eluted at a specific temperature. Maximum elution is reached at 40° C. C. iTRAQ results using two different oligos targeting *PHAROH* compared to PPIB reveal nucleolysin TIAR as the top hit. D. TIAR is pulled down by *PHAROH* oligos, and is specifically eluted at 40° C, but not by PPIB oligos. E. TIAR can be pulled down using additional oligos and in two different cell lines. RNase A treatment of the protein lysate diminishes TIAR binding to *PHAROH*, indicating that the interaction is RNA-dependent. F. Immunoprecipitation of TIAR enriches for *PHAROH* transcript, when compared to IgG and PPIB control.

For elution of *PHAROH*, we tested a range of temperatures and found that the elution efficiency reaches the maximum at 40° C, and thus we used this temperature for further experiments (Fig. 5B). The remaining level of *PHAROH* RNA on the beads was the direct inverse of the eluate (Fig. S5B). We chose *PPIB* as a negative control because it is a housekeeping mRNA that is expressed on the same order of magnitude as *PHAROH*, and is not expected to interact with the same proteins. We screened through ten oligos against *PPIB*, and found only one that pulled *PPIB* down at ∼60% efficiency, and eluted at the same temperature as *PHAROH* (Fig. S5C, S5D). Off-target RNA pulldown, such as *PHAROH* and *18S* rRNA, remained minimal when using the oligo antisense to *PPIB* (Fig. S5C).

To identify proteins that bind to *PHAROH*, we analyzed two independent oligos that target *PHAROH*, and two replicates of *PPIB*, on a single 4-plex iTRAQ (isobaric tag for relative and absolute quantitation) mass spectrometry cassette and identified a total of 690 proteins. By plotting the log_2_ enrichment ratio of *PHAROH* hits divided by *PPIB* hits, quadrant I will contain proteins that both oligos against *PHAROH* recognize, and quadrant III will be enriched for proteins that bind specifically to *PPIB*. Quadrant III was enriched for keratins, elongation factors, and ribosomal proteins. Interestingly, the top hit in quadrant I is nucleolysin TIAR (TIAL-1), an RNA-binding protein that controls mRNA translation by binding to AU-rich elements in the 3’ UTR of mRNA (Fig. 5C, Table 2) (Mazan-Mamczarz et al., 2006). TIAR is present in <10% of all experiments queried on Crapome.org (31/411). Immunoblots for TIAR confirm the mass spectrometry data in that TIAR is specific to *PHAROH* pull-down oligos, and also is eluted at 40° C (Fig 5D). Additional controls that are not complementary to the mouse genome and oligos targeting *PHAROH* also confirm the TIAR hit, and it is reproducible in two independent HCC cell lines (Fig. 5E). RNase A treatment of the lysate largely abolished the interaction, which indicates that the interaction is RNA mediated, and not the result of direct binding to the oligo (Fig. 5E). Immunoprecipitation of TIAR and subsequent extraction of interacting RNA shows enrichment for *PHAROH* when compared to *PPIB* and IgG control (Fig. 5F). Thus, together these data indicate that TIAR is a bona fide interactor of *PHAROH*.

**Table 2.**
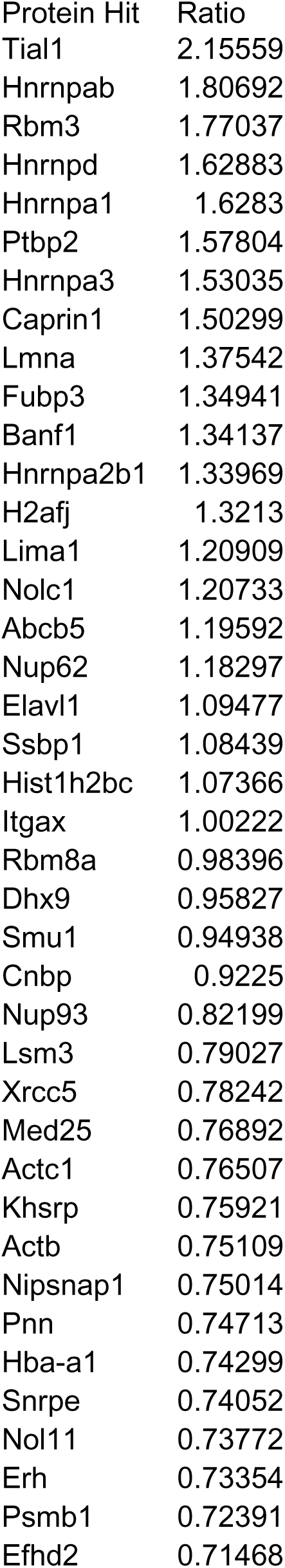
Top protein candidates that interact with PHAROH. Candidates with log2 fold change > 0.5 were used and the values from the two oligos were averaged and ranked from highest ratio of *PHAROH*/*Ppib* to lowest. TIAR (Tial1) is the top hit.

| Protein Hit | Ratio |
| --- | --- |
| Tial1 | 2.15559 |
| Hnrnpab | 1.80692 |
| Rbm3 | 1.77037 |
| Hnrnpd | 1.62883 |
| Hnrnpa1 | 1.6283 |
| Ptbp2 | 1.57804 |
| Hnrnpa3 | 1.53035 |
| Caprin1 | 1.50299 |
| Lmna | 1.37542 |
| Fubp3 | 1.34941 |
| Banf1 | 1.34137 |
| Hnrnpa2b1 | 1.33969 |
| H2afj | 1.3213 |
| Lima1 | 1.20909 |
| Nolc1 | 1.20733 |
| Abcb5 | 1.19592 |
| Nup62 | 1.18297 |
| Elavl1 | 1.09477 |
| Ssbp1 | 1.08439 |
| Hist1h2bc | 1.07366 |
| Itgax | 1.00222 |
| Rbm8a | 0.98396 |
| Dhx9 | 0.95827 |
| Smu1 | 0.94938 |
| Cnbp | 0.9225 |
| Nup93 | 0.82199 |
| Lsm3 | 0.79027 |
| Xrcc5 | 0.78242 |
| Med25 | 0.76892 |
| Actc1 | 0.76507 |
| Khsrp | 0.75921 |
| Actb | 0.75109 |
| Nipsnap1 | 0.75014 |
| Pnn | 0.74713 |
| Hba-a1 | 0.74299 |
| Snrpe | 0.74052 |
| Nol11 | 0.73772 |
| Erh | 0.73354 |
| Psmb1 | 0.72391 |
| Efhd2 | 0.71468 |

### A 71-nt sequence in PHAROH has four TIAR binding sites

A previous study on TIAR has mapped its RNA recognition motif across the transcriptome (Meyer et al., 2018). Analysis of *PHAROH*’s sequence reveals that TIAR binding sites are enriched in the 5’ end of the transcript of both isoforms (Fig. 6A). To determine if there are any conserved structure within *PHAROH* that mediates this interaction, RNA folding prediction algorithms, mFold and RNAfold, were used. The two strongest TIAR binding sequences (TTTT and ATTT/TTTA) were mapped onto ten outputted predicted structures (Fig. S6A). Strikingly, four out of the seven binding sites consistently mapped to a hairpin that was conserved throughout all predicted structures. Three of the strongest binding motifs localize to the stem of the hairpin, while one secondary motif resides in a bulge (Fig. 6B). These data indicate that the sequence is a highly concentrated site for TIAR binding, and is designed to potentially sequester multiple copies of TIAR.

**Figure 6.**
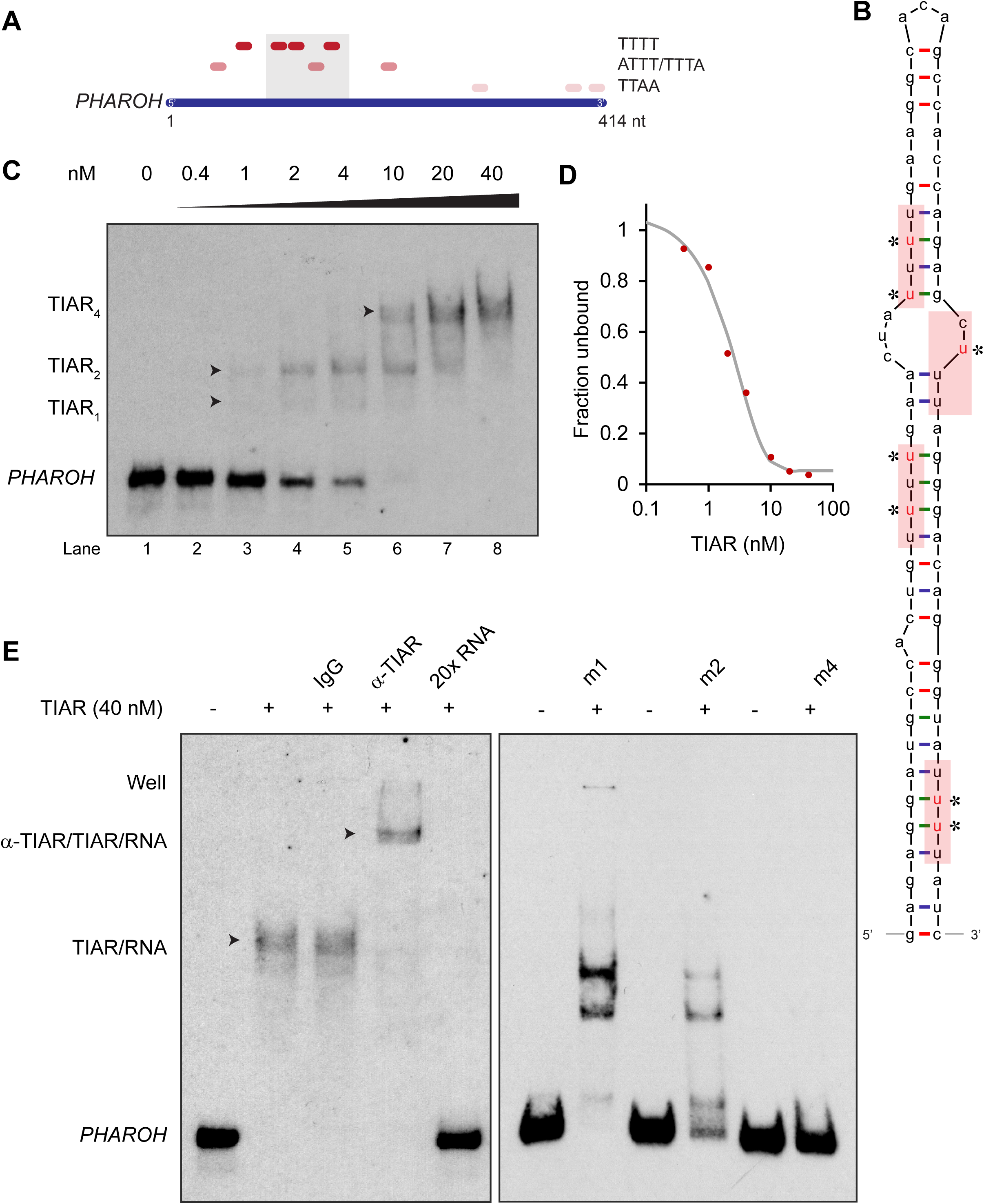
TIAR binds to the 5’ end of PHAROH. A. Sequence analysis of *PHAROH* with published TIAR binding motifs shows a preference for the 5’ end of *PHAROH*. B. Schematic of the conserved hairpin of *PHAROH* that contains four potential TIAR binding sites indicated in the red boxes. Mutations created within the *PHAROH* hairpin are indicated in red asterisks. C. RNA EMSA of the 71-nt *PHAROH* hairpin with human recombinant TIAR shows three sequential shifts as TIAR concentration increases. D. Densitometry analysis of the free unbound probe estimates the dissociation constant of TIAR as ∼2 nM. E. TIAR/PHAROH binding is specific as a supershift is created when adding antibody against TIAR, and the interaction can be competed out using 20x unlabeled RNA. RNA EMSA of the mutant hairpins reveals decreasing affinity for TIAR. M1 shows high signal of single and double occupancy forms, and m2 has reduced signal overall. When all four sites are mutated, binding is nearly abolished.

RNA electromobility shift assay (EMSA) of the hairpin and recombinant human TIAR showed that as TIAR concentration increases, it binds to the *PHAROH* hairpin multiple times (Fig. 6C). TIAR has a preference to bind two and four times, rather than once or three times. Densitometry quantification of the remaining free probe shows that TIAR has an approximate dissociation constant of 2 nM, consistent with the literature (Kim et al., 2011) (Fig. S6B). Addition of an antibody against TIAR creates a supershift, showing that the interaction is specific, while addition of IgG does not. The interaction can be abolished with addition of 20x unlabeled probe as well (Fig. 6E, left panel).

To determine if binding of TIAR is specific to the sequence and mapped motifs, we created sequential mutations of the hairpin by changing the non-canonical Watson-Crick base pairs (starred and in red) to canonical ones (Fig. 6B). Mutation of the first binding site (m1) slightly reduced specificity of TIAR to the hairpin, but changes the preference of TIAR binding to one and two units (Fig. 6E, right panel). Mutation of m2 greatly reduced TIAR association, and only two bands are highly visible (Fig. 6E, right panel). However, mutation of three binding sites (m3) did not appreciably change the pattern, as compared to m2, perhaps suggesting that the weaker binding site is only used cooperatively (Fig. S6C). Mutation of all four binding sites (m4) showed minimal TIAR binding (Fig. 6E). Taken together, these data indicate that TIAR binds directly to the 71-nt sequence on *PHAROH*, which can fold into a hairpin, and preferentially binds two or four times.

### PHAROH modulates c-Myc translation by sequestering TIAR

TIAR has been shown to bind to the 3’ UTR of mRNAs containing AU-rich elements in order to inhibit their translation (Mazan-Mamczarz et al., 2006). It has also been shown that TIAR binds to the 3’ UTR of *c-Myc* mRNA (Liao et al., 2007). Our data suggests that *PHAROH* serves to competitively sequester TIAR in order to allow for increased *c-MYC* translation. Thus, knockout or knockdown of *PHAROH* will free additional TIAR molecules to bind to the 3’ UTR of *c-Myc* and inhibit its translation.

We began by determining where TIAR binds to *c-Myc* mRNA. Mapping PAR-CLIP reads from (Meyer et al., 2018) shows two distinct binding sequences on the human *c-MYC* mRNA, but only one sequence maps to the mouse genome. The stretch of 53-nt sequence has three distinct regions that are enriched in poly-uridines, but structural prediction largely places the sequences in a loop formation (Fig. S7A, S7B). RNA EMSA of the 53-nt 3’ UTR and recombinant TIAR showed preference for a singular binding event, and three events are only seen when the binding reaction is saturated by TIAR (Fig. 7A). ASO mediated knockdown of *PHAROH* shows reduction of c-MYC protein similar to the knockouts, but no change in mRNA levels, or TIAR protein levels (Fig. 7B, S7C). While mRNAs are generally much more highly expressed than lncRNAs, *c-Myc* is only 3-fold more expressed than *PHAROH* in HCC cell lines (Fig. 7B). In addition, there are multiple TIAR binding sites on *PHAROH*, which increases the feasibility of a competition model (Fig. 7B).

**Figure 7.**
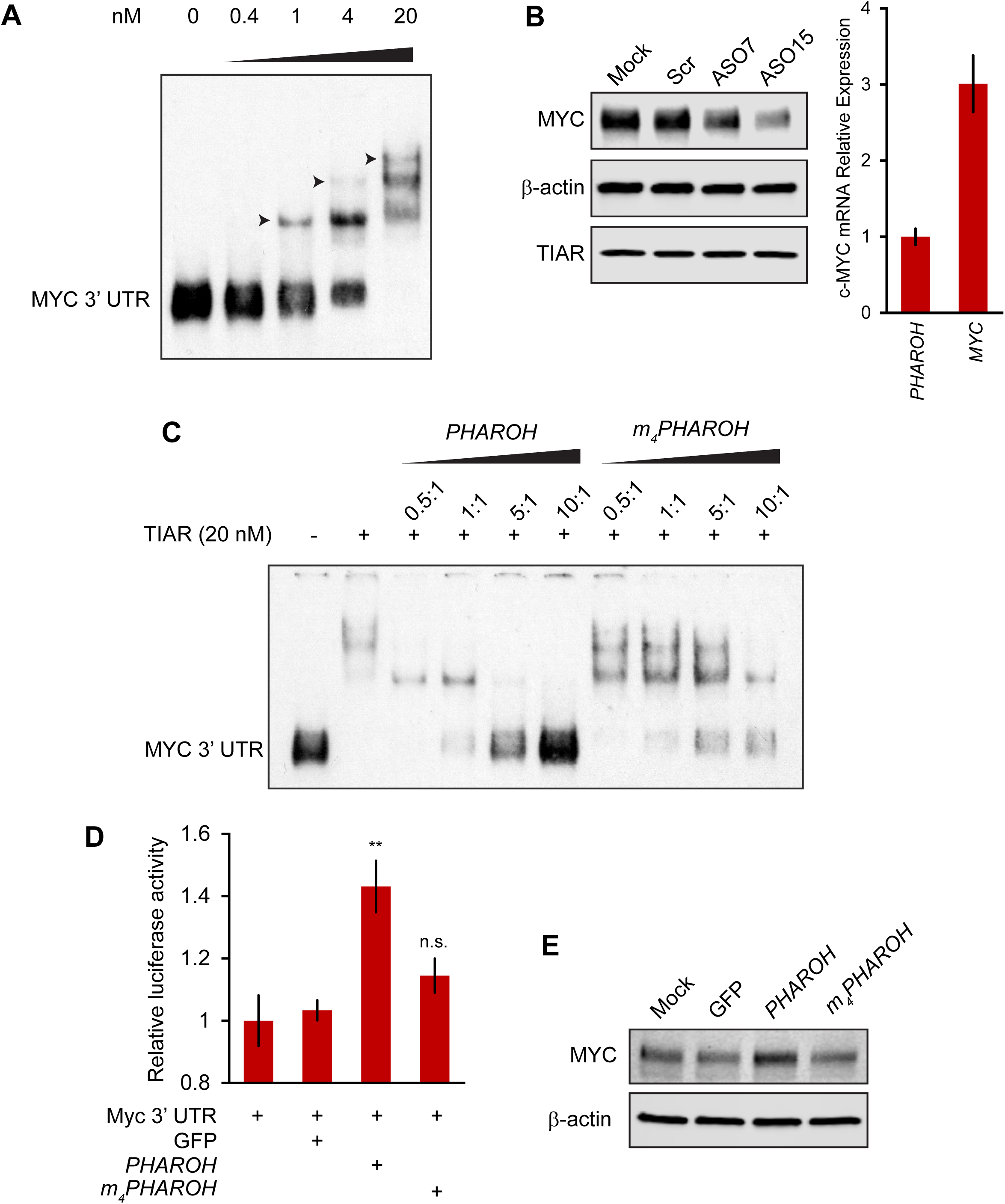
Loss of *PHAROH* releases TIAR, which inhibits c-Myc translation. A. RNA EMSA of the 53-nt c-Myc 3’ UTR fragment shows that TIAR has three potential binding sites, but prefers a single binding event (note arrows) B. Knockdown of *PHAROH* reduces c-MYC protein levels, but not TIAR levels, even though c-MYC is expressed 3-fold higher than *PHAROH*. C. Wildtype *PHAROH* hairpin is able to compete out the MYC-TIAR interaction, but the mutated hairpin is not as effective in competing with the Myc-TIAR interaction. D. Luciferase activity is increased with the addition of *PHAROH* but not with *m_4_PHAROH*. E. Overexpression of *PHAROH* increases c-MYC protein expression, but overexpression of *m_4_PHAROH* does not change c-MYC levels appreciably. **p < 0.01; Student’s t-test.

Next, we tested this hypothesis *in vitro*, by allowing TIAR to bind to the 53-nt c-Myc 3’ UTR, and titrating increasing amounts of *PHAROH* or the mutant *PHAROH* transcript. The wildtype *PHAROH* hairpin can be seen to compete with *c-Myc* very effectively at nearly all tested ratios, with near complete competition at 10:1 ratio (Fig. 7C). However, the fully mutant *PHAROH* was not able to compete with *c-Myc* nearly as effectively, and was only seen to be slightly effective at the 10:1 ratio (Fig. 7C). This data suggests that the *PHAROH* has the capability to successfully compete with the *c-Myc* 3’ UTR binding site in a sequence dependent manner.

In addition, we cloned the full length *c-Myc* 3’UTR into a dual luciferase reporter construct in order to test our hypothesis in cells. We found that addition of *PHAROH* does indeed increase the luciferase signal by ∼50% in a dose dependent manner while the mutant *PHAROH* did not (Fig. 7D, S7D).

Given that the knockdown or knockout of *PHAROH* reduces c-MYC levels due to the release of TIAR, we asked whether c-MYC protein levels would change in the context of *PHAROH* overexpression. Compared to GFP transfection, overexpression of *PHAROH* increases *c-MYC* protein levels; however, overexpression of mutant *PHAROH* did not change the protein levels of c-MYC (Fig. 7E). Modulation of *PHAROH* or TIAR levels did not have an effect on *c-Myc* mRNA levels (Fig. S7E).

## Discussion

Studies of the transcriptome have shed important insights into the potential role of the non-coding RNA portion of the genome in basic biology as well as disease. As such, lncRNAs can serve as biomarkers, tumor suppressors, or oncogenes, and have great potential as therapeutic targets (reviewed in Arun et al., 2018). Here, we identified a lncRNA, *PHAROH*, that is upregulated in mouse ESCs, embryonic and regenerating adult liver and in HCC. It also has a conserved human ortholog, which is upregulated in human patient samples from cirrhotic liver and HCC. Genetic knockout or ASO knockdown of *PHAROH* results in a reduction of cell proliferation, migration, and colony formation.

To elucidate the molecular mechanism through which *PHAROH* acts in proliferation, we used RNA-seq and mass spectrometry to provide evidence that *PHAROH* regulates c-MYC translation via sequestering the translational repressor TIAR in *trans*. Modulation of *PHAROH* levels reveal that it is positively correlated with c-MYC protein level, which is well known to be associated with HCC and is amplified in nearly 50% of HCC tumors (Peng et al., 1993). In addition, c-MYC has been characterized as a critical player in liver regeneration (Zheng et al., 2017). We identified TIAR as an intermediate player in the *PHAROH*-c-MYC axis, which has been reported to bind to the 3’ UTR of c-MYC mRNA and suppress its translation (Mazan-Mamczarz et al., 2006). While TIAR is an RNA-binding protein that is known for its role in stress granules (Kedersha et al., 1999), we do not detect stress granule formation in our HCC cell lines as assayed by immunofluorescence for TIAR (Fig. S7F). As such, the role of *PHAROH*-TIAR lies outside the context of stress granule function. Interestingly, overexpression of TIAR is a negative prognostic marker for HCC survival (Fig. S7G) (Uhlen et al., 2017). As the primary mutation of HCC is commonly amplification of c-MYC, it is possible that TIAR is upregulated in an attempt to curb c-MYC expression.

Our analysis maps the *PHAROH*-TIAR interaction to predominantly occur at a 71-nt hairpin at the 5’ end of *PHAROH*. While *PHAROH* has two main isoforms that are selectively expressed in ESCs and HCC, the hairpin is commonly expressed in both isoforms. TIAR has been classified as an ARE binding protein that recognizes U-rich and AU-rich sequences. Kinetic and affinity studies have found that TIAR has a dissociation constant of ∼1 nM for U-rich sequences, and ∼14 uM for AU-rich sequences (Kim et al., 2011). One question that is apparent in the RNA-binding protein field is how RBPs acquire their specificity. While there have been studies that analyze target RNA structure or RNA recognition motif structure, why RBPs bind one transcript over another with a similar sequence is still an open question. For example, the 3’ UTR of c-Myc contains multiple U-rich stretches, ranging from 3 to 9 resides. It has been reported that TIAR binds efficiently to uridylate residues of 3-11 length, yet PAR-CLIP data only reveals two binding events in the human *c-MYC* transcript (Kim et al., 2011). In addition, the 53-nt fragment that was assayed in this study contained potentially six TIAR binding sites, yet RNA EMSA analysis revealed a preference for a single binding event (Fig. 7A). One explanation is that *PHAROH*’s hairpin has uniquely spaced TIAR binding sites. Because the absolute affinity of TIAR to U-rich sequences is relatively high, one molecule may sterically block additional binding events. However, if the binding sites are properly spaced, binding events will be ordered and perhaps even cooperative. The average gap between binding sites in the *c-Myc* fragment is 2 nt, while it is 10 nt in the *PHAROH* hairpin, which allows more flexibility in spacing between each bound protein.

In addition, one aspect that was not explored was the requirement for the formation of the hairpin for TIAR binding. Previous studies used synthesized linear oligos as substrates to test the kinetics of these RBPs, and we also mutated the hairpin in a way such that structure is preserved. TIAR contains three RNA recognition motifs (RRM), which typically recognizes single stranded RNA. Therefore, binding of TIAR to the 71-nt sequence of *PHAROH* would require unwinding of the potential hairpin, which is energetically unfavorable. It is also known that TIAR’s RRM2 mainly mediates ssRNA polyU-binding, but its dsRNA binding capabilities have not been explored (Kim et al., 2013). There are examples where multiple RRMs in tandem can allow for higher RNA binding affinity and possibly sandwiching dsRNA, and thus it would be possible that TIAR binding to the multiple sites on the *PHAROH* hairpin is cooperative (Allain et al., 2000).

While TIAR may be *PHAROH*’s top interacting protein, it is unknown whether *PHAROH* is TIAR’s highest interacting RNA. This would depend on the relative abundances of each RNA species that has the potential to bind TIAR, and TIAR’s expression level. This seems to be cell type specific, as TIAR was initially studied in immune cells and was shown to predominantly translationally repress *Tnf-α* through binding of the AU-rich sequence in the 3’ UTR (Piecyk et al., 2000). In our cell lines, *Tnf-α* is not expressed. Conversely, a screen for proteins that bind to the *Tnf-α* 3’ UTR may not necessarily indicate TIAR as a binder, as evidenced by a recent study (Ma & Mayr, 2018). Another recent study had shown that lncRNA *MT1JP* functions as a tumor suppressor and had the capability to bind to TIAR, which suppresses the translation of p53 (Liu et al., 2016). However, *MT1JP* is largely cytoplasmic, while TIAR in our context is mainly nuclear. Thus, while TIAR may bind additional mRNAs or lncRNAs, it seems that one of the main targets in HCC cell lines is *c-Myc*, as supported by statistically significant promoter enrichment of the downstream targets.

In summary, we have identified a lncRNA, *PHAROH*, that is enriched in ESCs and dysregulated in HCC, and found that it acts to sequester TIAR through a hairpin structure in order to regulate *c-MYC* translation. Additionally, based on synteny and upregulation in human HCC samples, we identified *LINC00862* as the possible human ortholog of PHAROH (Fig. 1D). Future studies will reveal the therapeutic potential of targeting *PHAROH* to impact liver development/regeneration and HCC.

## Experimental procedures

### Cell culture and genomic PCR

All cell culture reagents were obtained from Gibco (Life Technologies), unless stated otherwise. Hepa1-6 (CRL-1830) and Hepa1c1c7 (CRL-2026) cells were obtained from ATCC. Both cell lines were maintained in DMEM supplemented with 10% FBS and 1% penicillin/streptomycin. Cells were cultured in a humidified incubator at 37° C and 5% CO_2_. Half-life of RNA was determined by adding α-amanitin to a final concentration of 5 µg/mL. Genomic DNA was isolated using DNeasy Blood & Tissue (Qiagen). Primers used are listed in the Table 3.

**Table 3.**
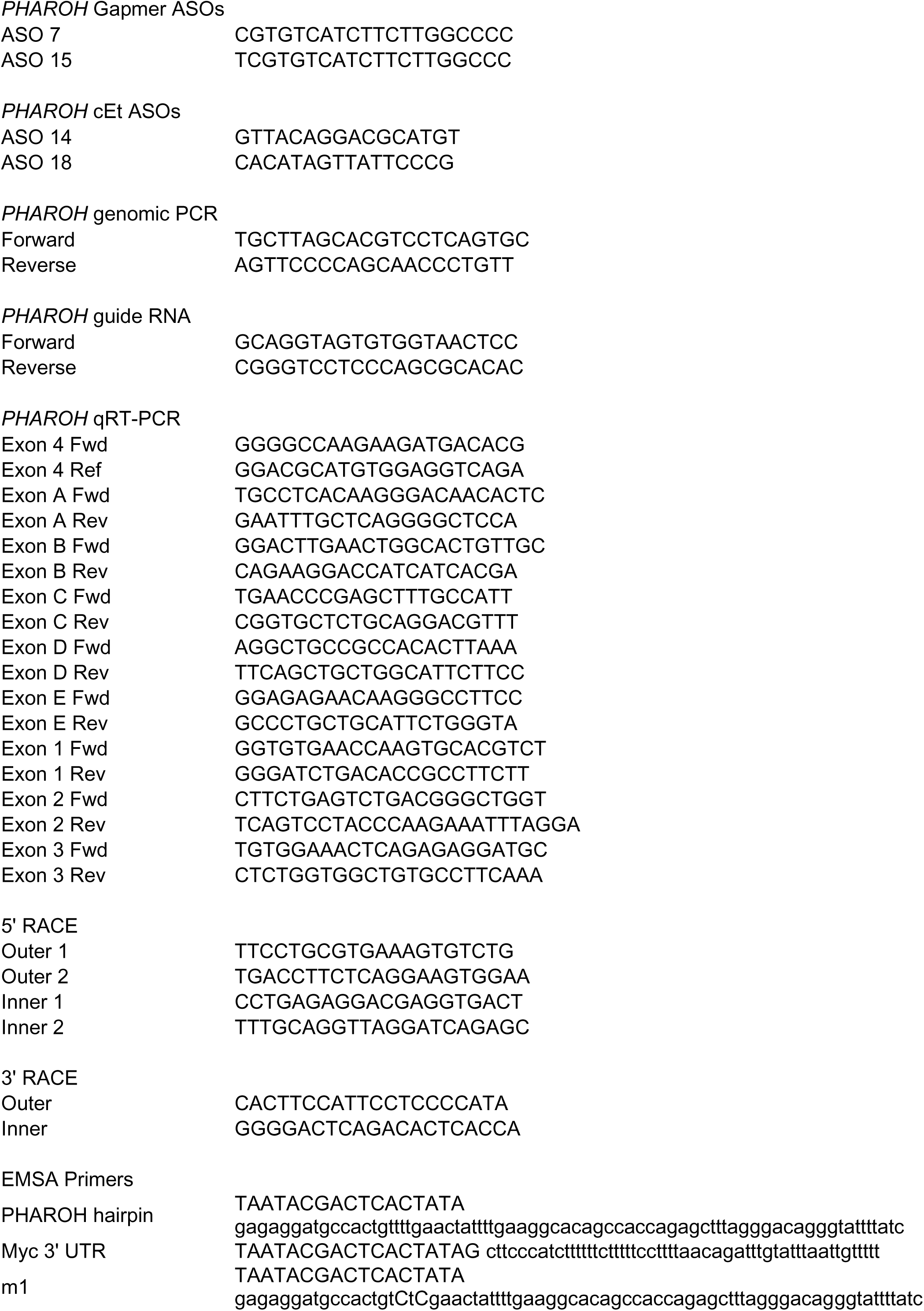
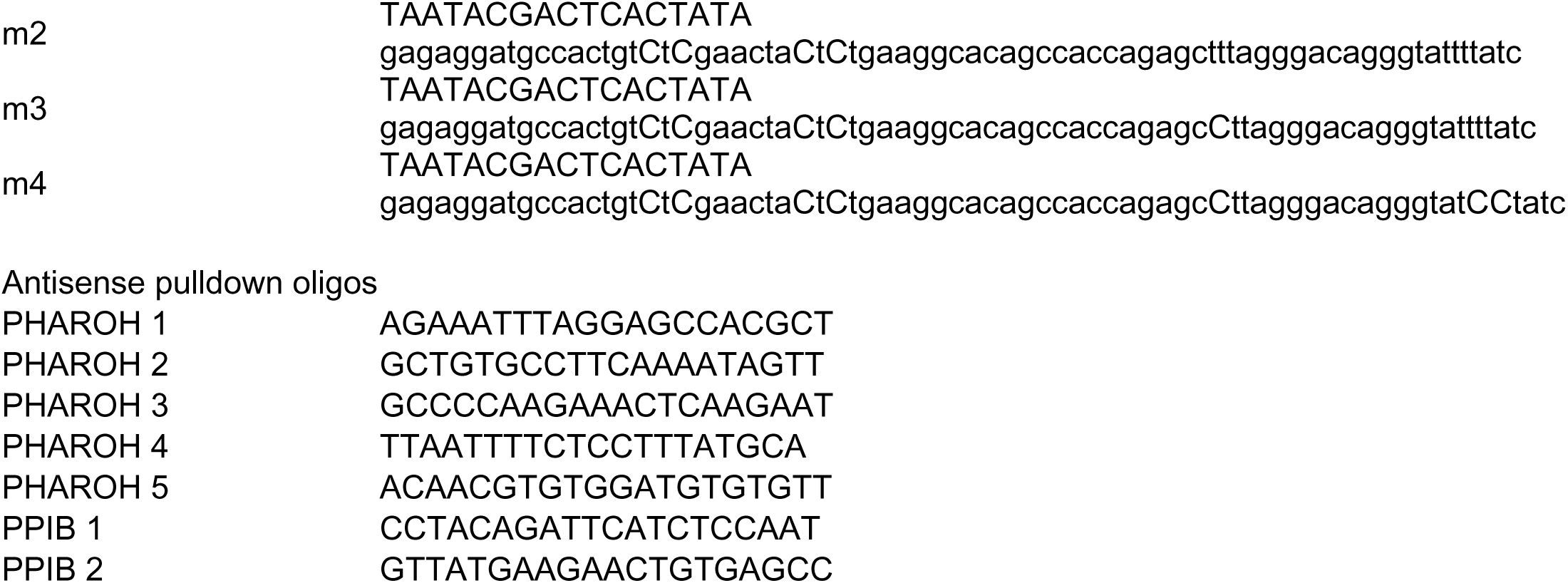
Sequences used in this study.

### Cellular Fractionation

Cellular fractionation was performed according to (https://link.springer.com/protocol/10.1007%2F978-1-4939-4035-6_1). In brief, cells were collected and resuspended in NP-40 lysis buffer. The cell suspension is overlaid on top of a sucrose buffer and centrifuged at 3,500 x g for 10 minutes to pellet the nuclei. The supernatant (cytoplasm) is collected and the nuclei are resuspended in glycerol buffer and urea buffer is added to separate the nucleoplasm and chromatin. The cells are centrifuged at 14,000 x g for 2 minutes and the supernatant (nucleoplasm) is collected, while the chromatin-RNA is pelleted.

### DEN administration

Mice were injected intraperitoneally with diethylnitrosamine (DEN) at 14 days of age as described (Garcia-Irigoyen et al., 2015). DEN-treated mice, and the corresponding controls injected with saline, were sacrificed at 5, 8, and 11 months post injection.

### Partial hepatectomy

Two-thirds partial hepatectomy (PH) and control sham operations (SH) were performed as reported (Berasain et al., 2005). Two SH and four PH mice were sacrificed at 3, 6, 24, 48 and 72 hours after surgery.

### Human samples

Samples from patients included in the study were provided by the Biobank of the University of Navarra and were processed following standard operating procedures approved by the Ethical and Scientific Committees. Liver samples from healthy patients were collected from individuals with normal or minimal changes in the liver at surgery of digestive tumors or from percutaneous liver biopsy performed because of mild alterations of liver function. Samples for cirrhotic liver and HCC were obtained from patients undergoing partial hepatectomy and/or liver transplantation

### Immunoblotting

To determine protein levels in our system, we used 10% SDS-PAGE gels. Gels were loaded with 1-μg protein per well (Bradford assay). The following antibodies were used: β-actin (1:15,000; Sigma), c-Myc (1:1000; CST), TIAR (1:1000; Cell Signaling). IRDye-800CW was used as a fluor for secondary anti-rabbit antibodies, and IRDye-680RD was used for mouse secondary antibodies. Blots were scanned using the Li-Cor Odyssey Classic.

### Immunoprecipitation

For TIAR immunoprecipitation, one 10 cm plate of Hepa1c1c7 cells at 80% confluence was lysed in 1 mL Pierce IP Lysis Buffer (supplemented with 100 U/mL SUPERase-IN and 1X Roche protease inhibitor cocktail) and incubated on ice for 10 min. Lysates centrifuged at 13,000xg for 10 minutes. 3 ug of TIAR antibody or rabbit IgG were incubated with the lysate at 4°C for 1 hour. 16 uL of Protein A magnetic beads were washed and added to the lysate and incubated for an additional 30 minutes at 4°C. 50% of beads were resuspended in Laemmli buffer for western blotting and RNA was isolated from the remaining beads using TRIzol.

### Immunofluorescence staining

#1.5 round glass coverslips were prepared by acid-cleaning prior to seeding cells. Staining was performed as published previously (Spector, D.L. and H.C. Smith. 1986. Exp. Cell Res. 163, 87-94). In brief, cells were fixed in 2% PFA for 15 min, washed with PBS, and permeabilized in 0.2% Triton X-100 plus 1% normal goat serum (NGS). Cells were washed again in PBS+1% NGS, and incubated with TIAR antibody (1:2000; CST) for 1 hour at room temp in a humidified chamber. Cells were washed again PBS+1% NGS, and incubated with Goat anti-Rabbit IgG (H+L) Cross-Adsorbed Secondary Antibody, Alexa Fluor 488 (1:1000; Thermo Fisher) secondary antibody for 1 hour at room temp. Cover slips were washed with PBS before mounting with ProLong Diamond antifade (Thermo Fisher).

### Cell viability assays

Cells were seeded at a density of 10,000 cells/well (100 µl per well) into 24-well plates and treated with 2.5 µM of either a *PHAROH*-specific ASO or scASO. Cells were grown for 96 h at 37°C. 10 µl MTT solution (Cell Growth Determination Kit, MTT based; Sigma) was added to the wells and incubated for 4 h at 37°C. Next, 100 µl MTT solvent was added directly to the wells to ensure total solubility of the formazan crystals and incubated for 10 min with shaking. Measurements of absorbance at 570 nm were performed using a SpectraMax i3 Multi-Mode Detection Platform (Molecular Devices). Background absorbance at 690 nm was subtracted. Cells were also trypsinized, pelleted, and manually counting using a hemocytometer.

### RNA antisense pulldown and mass spectrometry

RNA antisense pulldown—Cells were lysed on a 10 cm plate in 1 mL IP lysis buffer (IPLB, 25 mM Tris-HCl pH 7.4, 150 mM NaCl, 1% NP-40, 1 mM EDTA, 5% glycerol, supplemented with 100 U/mL SUPERase-IN and 1X Roche protease inhibitor cocktail) for 10 minutes, and lysate was centrifuged at 13,000xg for 10 minutes. Cell lysate was adjusted to 0.3 mg/mL (Bradford assay). 100 pmol of biotinylated oligo was added to 500 uL of lysate and incubated at room temperature for 1 hour with rotation. 100 uL streptavidin Dynabeads were washed in IPLB, added to the lysate, and incubated for 30 minutes at room temperature with rotation. Beads were washed three times with 1 mL lysis buffer. For determining temperature for optimal elution, beads were then resuspended in 240 uL of 100 mM TEAB and aliquoted into eight PCR tubes. Temperature was set on a veriflex PCR block and incubated for 10 minutes. Beads were captured and TRIzol was added to the eluate and beads. Once optimal temperature is established, the beads were resuspended in 90 uL of 100 mM TEAB, and incubated at 40° C for 10 minutes. TRIzol was added to 30 uL of the eluate, another 30 uL was kept for western blots, and the last 30 uL aliquot was sent directly for mass spectrometry. Oligo sequences available in Table 3.

Tryptic digestion and iTRAQ labeling—Eluted samples were reduced and alkylated with 5 mM DTT and 10 mM iodoacetamide for 30 min at 55 °C, then digested overnight at 37 °C with 1 μg Lys-C (Promega, VA1170) and dried in vacuo. Peptides were then reconstituted in 50 μl of 0.5 M TEAB/70% ethanol and labeled with 4-plex iTRAQ reagent for 1 h at room temperature essentially as previously described (Ross et al., 2004). Labeled samples were then acidified to <pH 4 using formic acid, combined and concentrated in vacuo until ∼10 μl remained.

Two-dimensional fractionation—Peptides were fractionated using a Pierce™ High pH Reversed-Phase Peptide Fractionation Kit (Thermo Scientific, 84868) according to the manufacturer’s instructions with slight modifications. Briefly, peptides were reconstituted in 150 μl of 0.1% TFA, loaded onto the spin column, and centrifuged at 3000 × g for 2 min. Column was washed with water, and then peptides were eluted with the following percentages of acetonitrile (ACN) in 0.1% triethylamine (TEA): 5%, 7.5%, 10%, 12.5%, 15%, 20%, 30%, and 50%. Each of the 8 fractions was then separately injected into the mass spectrometer using capillary reverse-phase LC at low pH.

Mass spectrometry—An Orbitrap Fusion Lumos mass spectrometer (Thermo Scientific), equipped with a nano-ion spray source was coupled to an EASY-nLC 1200 system (Thermo Scientific). The LC system was configured with a self-pack PicoFrit™ 75-μm analytical column with an 8-μm emitter (New Objective, Woburn, MA) packed to 25 cm with ReproSil-Pur C18-AQ, 1.9 μM material (Dr. Maish GmbH). Mobile phase A consisted of 2% acetonitrile; 0.1% formic acid and mobile phase B consisted of 90% acetonitrile; 0.1% formic acid. Peptides were then separated using the following steps: at a flow rate of 200 nl/min: 2% B to 6% B over 1 min, 6% B to 30% B over 84 min, 30% B to 60% B over 9 min, 60% B to 90% B over 1 min, held at 90% B for 5 min, 90% B to 50% B over 1 min and then flow rate was increased to 500 μl/min as 50% B was held for 9 min. Eluted peptides were directly electrosprayed into the MS with the application of a distal 2.3 kV spray voltage and a capillary temperature of 300 °C. Full-scan mass spectra (Res = 60,000; 400–1600 m/z) were followed by MS/MS using the “Top Speed” method for selection. High-energy collisional dissociation (HCD) was used with the normalized collision energy set to 35 for fragmentation, the isolation width set to 1.2 and a duration of 15 s was set for the dynamic exclusion with an exclusion mass width of 10 ppm. We used monoisotopic precursor selection for charge states 2+ and greater, and all data were acquired in profile mode.

### Database searching

Peaklist files were generated by Proteome Discoverer version 2.2.0.388 (Thermo Scientific). Protein identification was carried out using both Sequest HT (Eng et al., 1994) and Mascot 2.5 (Perkins et al., 1999) against the UniProt mouse reference proteome (57,220 sequences; 26,386,881 residues). Carbamidomethylation of cysteine, iTRplex (K), and iTRplex (N-term) were set as fixed modifications, methionine oxidation, and deamidation (NQ) were set as variable modifications. Lys-C was used as a cleavage enzyme with one missed cleavage allowed. Mass tolerance was set at 20 ppm for intact peptide mass and 0.3 Da for fragment ions. Search results were rescored to give a final 1% FDR using a randomized version of the same Uniprot mouse database, with two peptide sequence matches (PSMs) required. iTRAQ ratio calculations were performed using Unique and Razor peptide categories in Proteome Discoverer.

### RNA Electromobility shift assay

DNA template used for in vitro synthesis of RNA probes were from annealed oligos. A T7 RNA polymerase promoter sequence was added to allow for in vitro transcription using the MEGAscript T7 transcription kit (Thermo Fisher). RNA was end labelled at the 3’ end with biotin using the Pierce RNA 3’ End Biotinylation Kit (Thermo Fisher). RNA quantity was assayed by running an RNA 6000 Nano chip on a 2100 Bioanalyzer. Six percent acrylamide gels (39:1 acrylamide:bis) (Bio-Rad) containing 0.5 X TBE were used for all EMSA experiments. Recombinant human TIAR (Proteintech) was added at indicated concentrations to the probe (∼2 fmol) in 20 uL binding buffer, consisting of 10 mM HEPES (pH 7.3), 20 mM KCL, 1 mM Mg_2_Cl_2_, 1 mM DTT, 30 ng/uL BSA, 0.01% NP-40, and 5% glycerol. After incubation at room temperature for 30 minutes, 10 uL of the samples were loaded and run for 1 hr at 100 V. The nucleic acids were then transferred onto a positively charged nylon membrane (Amersham Hybond-N+) in 0.5 X TBE for 30 minutes at 40 mAh. Membranes were crosslinked using a 254 nM bulb at 120 mJ/cm^2^ in a Stratalinker 1800. Detection of the biotinylated probe was done using the Chemiluminescent Nucleic Acid Detection Module Kit (Thermo Fisher 89880).

### 3’ UTR luciferase assay

The full length 3’ UTR of c-Myc was cloned into the pmirGLO Dual-Luciferase miRNA target expression vector (Promega). Luciferase activity was assayed in transfected cells using the Dual-Luciferase Reporter Assay (Promega). To evaluate the interaction between *PHAROH*, 3’ UTR of c-Myc, and TIAR, cells were transfected with the respective constructs using Lipofectamine 3000. Twenty-four hours later, firefly and Renilla luciferase activity was measured, and Renilla activity was used to normalize firefly activity.

### Single Molecule RNA FISH

#1.5 round glass coverslips were prepared by acid-cleaning and layered with gelatin for 20 minutes, prior to seeding MEF feeder cells and ESCs. Cells were fixed for 30 minutes in freshly-prepared 4% PFA (Electron Microscopy Sciences), diluted in D-PBS without CaCl_2_ and MgCl_2_ (Gibco, Life Technologies) and passed through a 0.45 µm sterile filter. Fixed cells were dehydrated and rehydrated through an ethanol gradient (50% - 75% - 100% - 75% - 50%- PBS) prior to permeabilization for 5 minutes in 0.5% Triton X-100. Protease QS treatment was performed at a 1:8,000 dilution. QuantiGene ViewRNA (Affymetrix) probe hybridizations were performed at 40°C in a gravity convection incubator (Precision Scientific), and incubation time of the pre-amplifier was extended to 2 hours. Nuclei were counter-stained with DAPI and coverslips mounted in Prolong Gold anti-face medium (www.spectorlab.labsites.cshl.edu/protocols).

Coverslips were imaged on a DeltaVision Core system (Applied Precision), based on an inverted IX-71 microscope stand (Olympus) equipped with a 60x U-PlanApo 1.40 NA oil immersion lens (Olympus). Images were captured at 1×1 binning using a CoolSNAP HQ CCD camera (Photometric) as z-stacks with a 0.2 µm spacing. Stage, shutter and exposure were controlled through SoftWorx (Applied Precision). Image deconvolution was performed in SoftWorx.

A spinning-disc confocal system (UltraVIEW Vox; PerkinElmer) using a scanning unit (CSU-X1; Yokogawa Corporation of America) and a charge-coupled device camera (ORCA-R2; Hamamatsu Photonics) fitted to an inverted microscope (Nikon) equipped with a motorized piezoelectric stage (Applied Scientific Instrumentation). Image acquisition was performed using Volocity versions 5 and 6 (PerkinElmer). Routine imaging performed using Plan Apochromat 60 or 100× oil immersion objectives, NA 1.4.

### RNA sequencing and analysis

Total RNA was isolated either directly from cryosections of the tumor tissue or from organotypic epithelial cultures using TRIzol according to the manufacturer’s instructions. RNA quality was assayed by running an RNA 6000 Nano chip on a 2100 Bioanalyzer. For high-throughput sequencing, RNA samples were required to have an RNA integrity number (RIN) 9 or above. TruSeq (Illumina) libraries for poly(A)+ RNA-seq were prepared from 0.5–1mg RNA per sample. To ensure efficient cluster generation, an additional gel purification step of the libraries was applied. The libraries were multiplexed (12 libraries per lane) and sequenced single-end 75 bp on the NextSeq500 platform (Illumina), resulting in an average 40 Million reads per library. Analysis was performed in GalaxyProject. In brief, reads were first checked for quality using FastQC (http://www.bioinformatics.babraham.ac.uk/projects/fastqc/), and a minimum Phred score of 30 was required. Reads were then mapped to the mouse mm10 genome using STAR (Dobin et al., 2013), and counts were generating using htseq-counts with the appropriate GENCODE M20 annotation. Deseq2 was then used to generate the list of differentially expressed genes (Love et al., 2014). Motif analysis was performed using HOMER (Heinz et al., 2010).

### Coding analysis

cDNA sequences of *PHAROH* and *GAPDH* were inputted into CPAT (http://lilab.research.bcm.edu/cpat/) or CPC (http://cpc.cbi.pku.edu.cn/programs/run_cpc.jsp) for analysis. PhyloCSF analysis was performed using the UCSC Genome Browser track hub (https://data.broadinstitute.org/compbio1/PhyloCSFtracks/trackHub/hub.DOC.html).

### Plasmid construction

eSpCas9(1.1) was purchased from Addgene (#71814). eSpCas9-2A-GFP was constructed by subcloning 2A-GFP from pSpCas9(BB)-2A-GFP (PX458) (Addgene #48138) into eSpCas9 using EcoRI sites. To construct eSpCas9-2A-mCherry, 2A-mCherry was amplified from mCherry-Pol II (Zhao et al., 2011), and an internal BbsI site was silently mutated. The PCR product was then cloned into eSpCas9 using EcoRI sites. The *PHAROH* construct was amplified using Hepa1-6 cDNA as a template and cloned into pCMV6 using BamHI and FseI. Mutant *PHAROH* was constructed by amplifying tiled oligos and cloned into pCMV6 using BamHI and FseI.

### CRISPR/Cas9 genetic knockout

To generate a genetic knockout of *PHAROH*, two sgRNAs targeting the promoter region were combined, creating a deletion including the TSS. Guide design was performed on Benchling (https://benchling.com) taking into account both off-target scores and on-target scores. The sgRNA targeting the gene body of *PHAROH* was cloned into a pSpCas9(BB)-2A-GFP vector (PX458, Addgene plasmid #48138) and the sgRNA targeting the upstream promoter region was cloned into a pSpCas9(BB)- 2A-mCherry vector. Hepa1-6 were transfected with both plasmids using the 4D-Nucleofector™ System (Lonza) using the EH-100 program in SF buffer. To select for cells expressing both gRNAs, GFP and mCherry double positive cells were sorted 48 hours post transfection, as single cell deposition into 96-well plates using a FACS Aria (SORP) Cell Sorter (BD). Each single cell clone was propagated and analyzed by genomic PCR and qRT-PCR to select for homozygous knockout clones. Cells transfected with a sgRNA targeting Renilla luciferase were used as a negative control. Sequences for sgRNAs and primers are provided in Table 3.

### Cell cycle analysis

Hoechst 33342 (Sigma) was added to cells at a final concentration of 5 μg/mL and incubated at 37° C for 1 hour. Cells were trypsinized and collected into a flow cytometry compatible tube. Profiles were analyzed using a FACS Aria (SORP) Cell Sorter (BD), gated according to DNA content and cell cycle phase, and sorted into Eppendorf tubes for subsequent RNA extraction and qRT-PCR analysis.

### Nucleofection

For transfection of ASOs using nucleofection technology (Lonza), ESCs were harvested following soaking off of feeder cells for one hour, washed in D-PBS (Gibco, Life Technologies) and passed through a 70 µm nylon cell strainer (Corning). Cell count and viability was determined by trypan blue staining on a Countess automated cell counter (Life Technologies). For each reaction, 3×10^6^ viable cells were resuspended in P3 Primary Cell solution (Lonza), mixed with 2 µM control or 2 µM target-specific ASO and transferred to nucleocuvettes for nucleofection on a 4D-Nucleofector System (Lonza) using program code “DC-100”. For plasmid nucleofections, 20 ug of plasmid was used and nucleofected using program code “CG-104”. Cells were subsequently transferred onto gelatinized cell culture plates containing pre-warmed and supplemented growth medium. Growth medium was changed once after 16 hours.

### Colony Formation Assay

200 Hepa1-6 cells were seeded in a 6-well plate. ASOs were added at the time of seeding at the indicated concentrations. Two weeks later, cells were fixed, stained with Giemsa, counted and photographed.

### 2’-O-Methoxyethyl (MOE) antisense oligonucleotides and knockdown analysis

Synthesis and purification of all 2’-MOE modified oligonucleotides was performed as previously described (Meng et al. 2014) by Ionis Pharmaceuticals. These ASOs are 20-mer oligonucleotides containing a phosphorothioate backbone, 2’-O-methoxyethyl modifications on the first and last five nucleotides and a stretch of ten DNA bases in the center. Constrained ethyl oligos are 16-mer oligonucleotides that contain modifications on the first and last 3 nucleotides and a stretch of ten DNA bases in the center. ASO sequences available in table 3.

### qRT-PCR

To assess knockdown efficiency TRIzol-extracted RNA was treated with RNAse-free DNAseI (Life Technologies) and subsequently reverse-transcribed into cDNA using TaqMan Reverse Transcription reagents and random hexamer oligonucleotides (Life Technologies). Real-time PCR reactions were prepared using Power SYBR Green Master Mix (Life Technologies) and performed on an ABI StepOnePlus Real-Time PCR system (Life Technologies) for 40 cycles of denaturation at 95°C for 15 seconds followed by annealing and extension at 60°C for 60 seconds. Primers were designed to anneal within an exon to detect both primary and processed transcripts. Primer specificity was monitored by melting curve analysis. For each sample, relative abundance was normalized to the housekeeping gene *PPIB* mRNA levels.

## Author Contributions

Conceptualization, A.T.Y. and D.L.S.; Methodology, A.T.Y., C.B., K.R., B.L., D.J.P., and D.L.S; Formal Analysis, A.T.Y, C.B., S.B., K.R., and D.J.P.; Investigation, A.T.Y, C.B., S.B., and K.R; Writing, A.T.Y. and D.L.S. produced the initial manuscript, and all others reviewed and commented the manuscript; Resources, C.B. and F.R; Supervision, D.L.S; Funding Acquisition, A.T.Y and D.L.S.

## Acknowledgments

We thank members of the Spector lab for critical discussions and advice throughout the course of this study. We would also like to thank the CSHL Cancer Center Shared Resources (Microscopy, Mass Spectrometry, Flow Cytometry, and Next-Gen Sequencing) for services and technical expertise (NCI 2P3OCA45508). This research was supported by NCI 5P01CA013106-Project 3 (D.L.S.) and NCI 5F31CA220997-02.

---

**Supplemental Figure 1.**
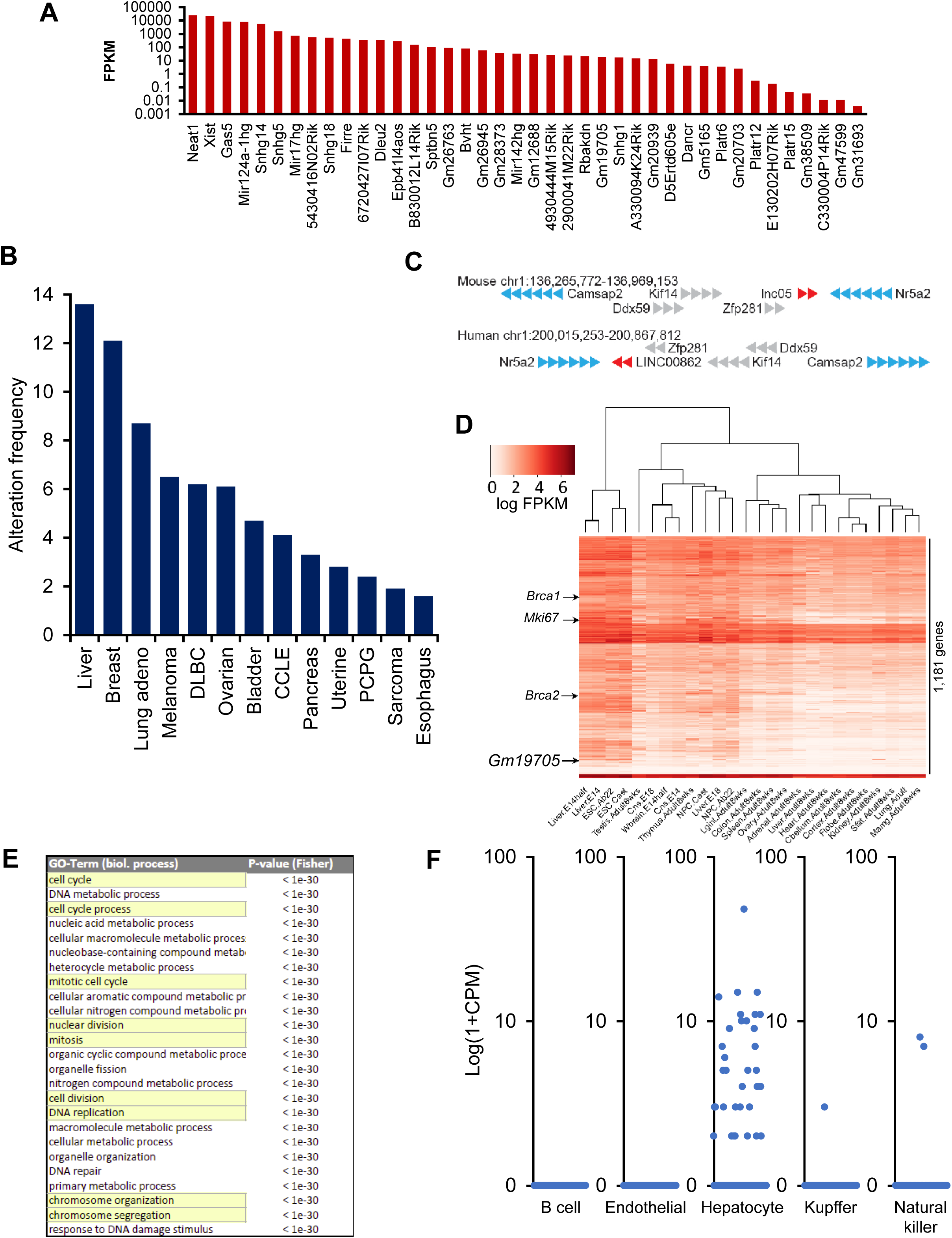
A. LncRNA screen identifies candidates with varying levels of expression in ESCs. B. *LINC00862* is altered in 13% of all HCC patient cases according to TCGA data. C. *Gm19705* gene locus on chromosome 1 shows that the order of the genes is conserved between mouse and human, but the order is reversed, suggesting a reversal event occurrence. D. Weighted gene correlation network analysis of *Gm19705* places it in a module with cell cycle genes and proliferation genes, such as *Brca1/2*, and *Mki67*. E. GO term analysis of the module containing *Gm19705* shows enrichment of genes related to cell cycle, mitosis, and DNA replication. F. Re-analysis of single cell data of adult liver (Tabula Muris et al., 2018) reveals expression of *Gm19705* is highly enriched in hepatocytes, but only a subset of the cells.

**Supplemental Figure 2.**
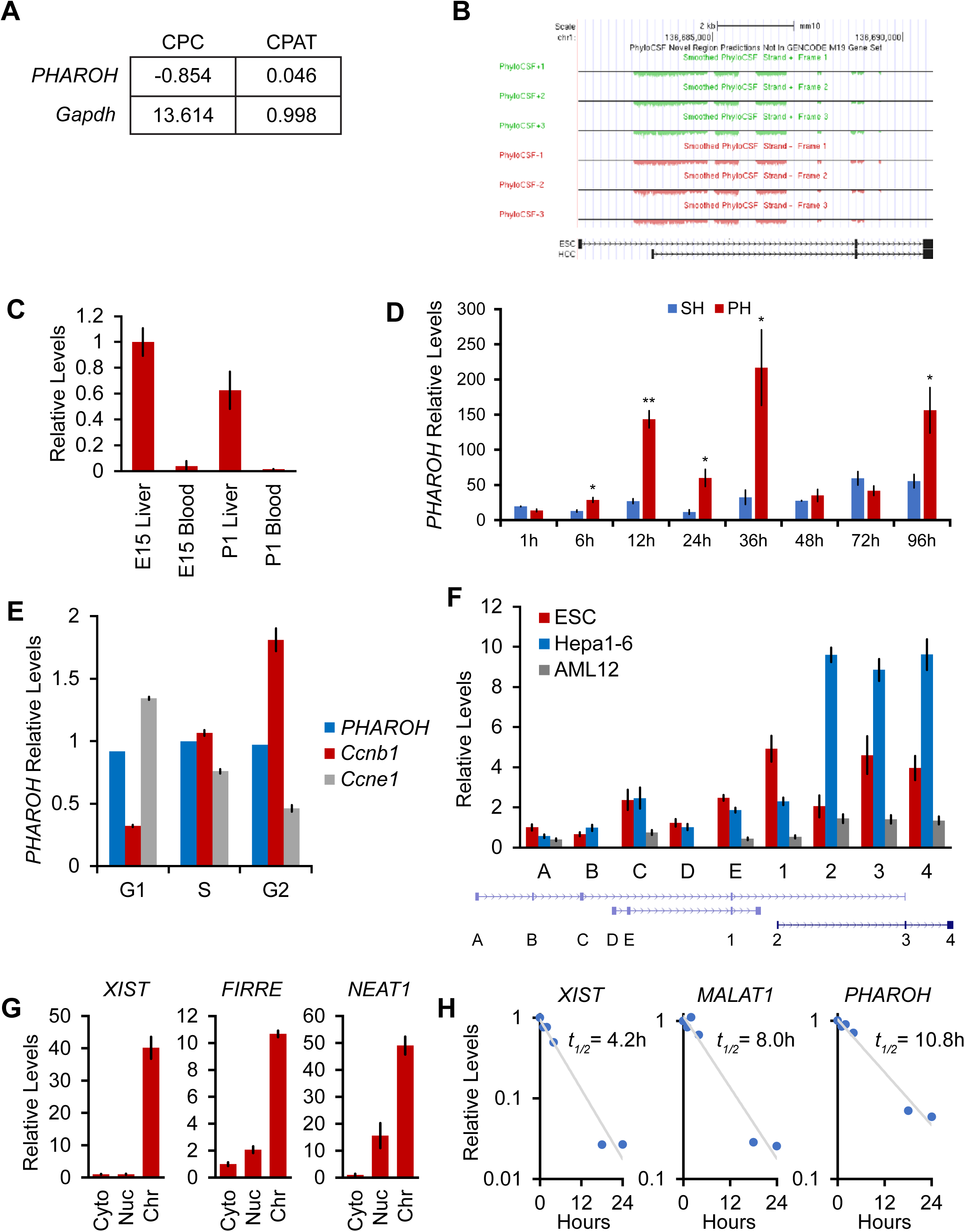
A. CPC and CPAT coding potential analysis for *PHAROH* and *Gapdh*. B. PhyloCSF tracks showing low coding potential for the *PHAROH* locus C. *PHAROH* is expressed in fetal liver, but not in the blood. D. Sham hepatectomy (SH) or partial hepatectomy (PH) of the liver, a model of liver regeneration, shows upregulation of *PHAROH* during time points of concerted cell division. *p < 0.05; **p < 0.01; ***p < 0.005; Student’s t-test. E. HepA1-6 cells were stained with Hoechst 33258 and sorted according to their cell cycle phase. qRT-PCR analysis shows that *PHAROH* does not cycle with the cell cycle, unlike *Ccnb1* and *Ccne1*. F. qRT-PCR of each annotated exon. Exons 1-4 are confirmed RACE exons. Isoform with exons 1, 3, and 4 is ESC specific, and the isoform with exons 2-4 is HCC specific. G. *XIST*, *FIRRE*, and *NEAT1* serve as additional controls for the cellular fractionation. H. Calculated RNA half-life based upon α-amanitin treated cells. *PHAROH* has a half-life of 10.8h, longer than that of *XIST* and *MALAT1*.

**Supplemental Figure 3.**
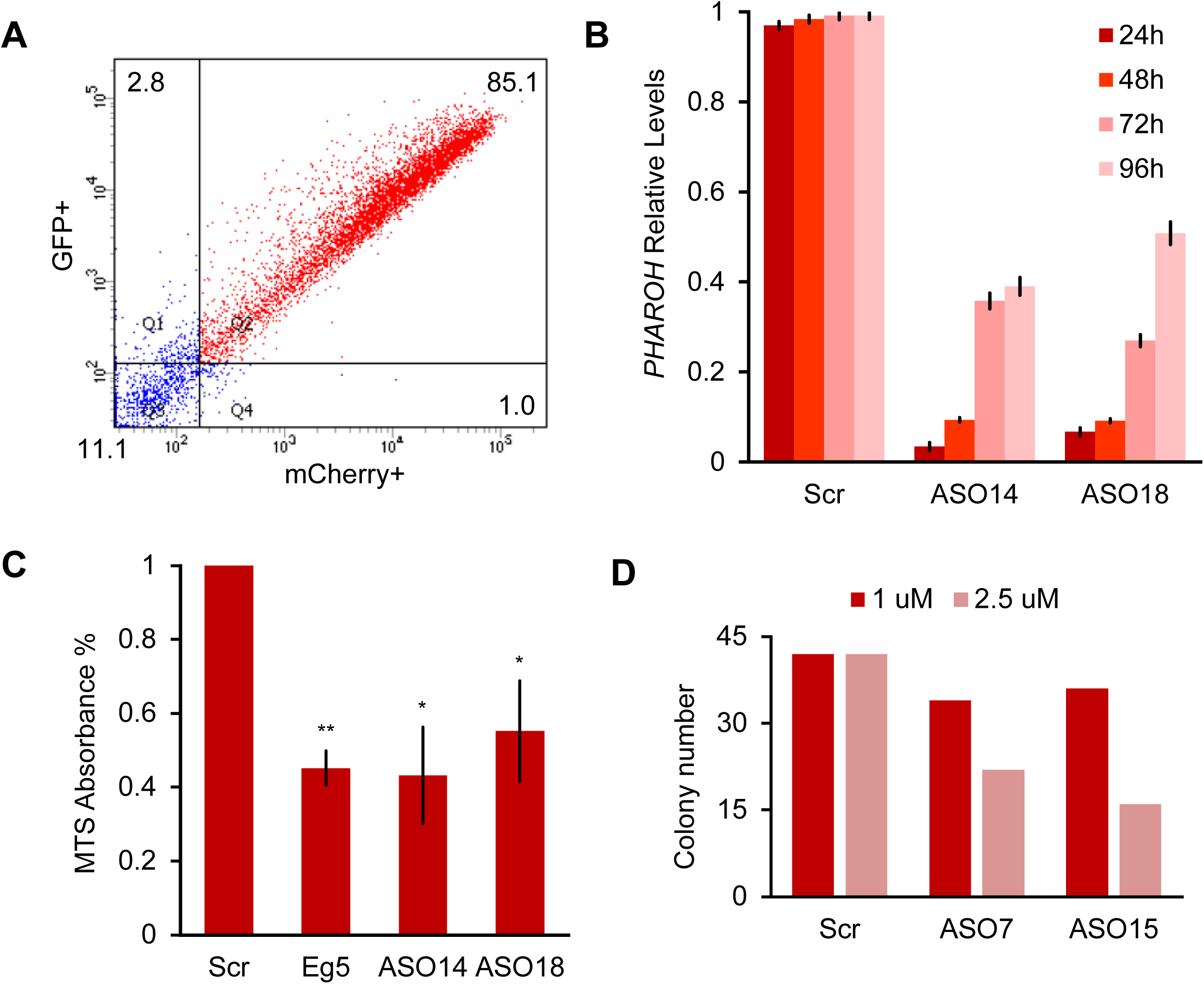
A. FACS for double GFP+/mCherry+ cells shows an 85.1% nucleofection efficiency for both plasmids. B. Knockdown of *PHAROH* using nucleofection of 2 μM ASO is effective over 96h. C. MTS assay for proliferation 96h after nucleofection. MTS absorbance is reduced by 50% in ASO treated samples targeting *PHAROH* and *Eg5*. D. Reduction of colony formation number is dose dependent.

**Supplemental Figure 4.**
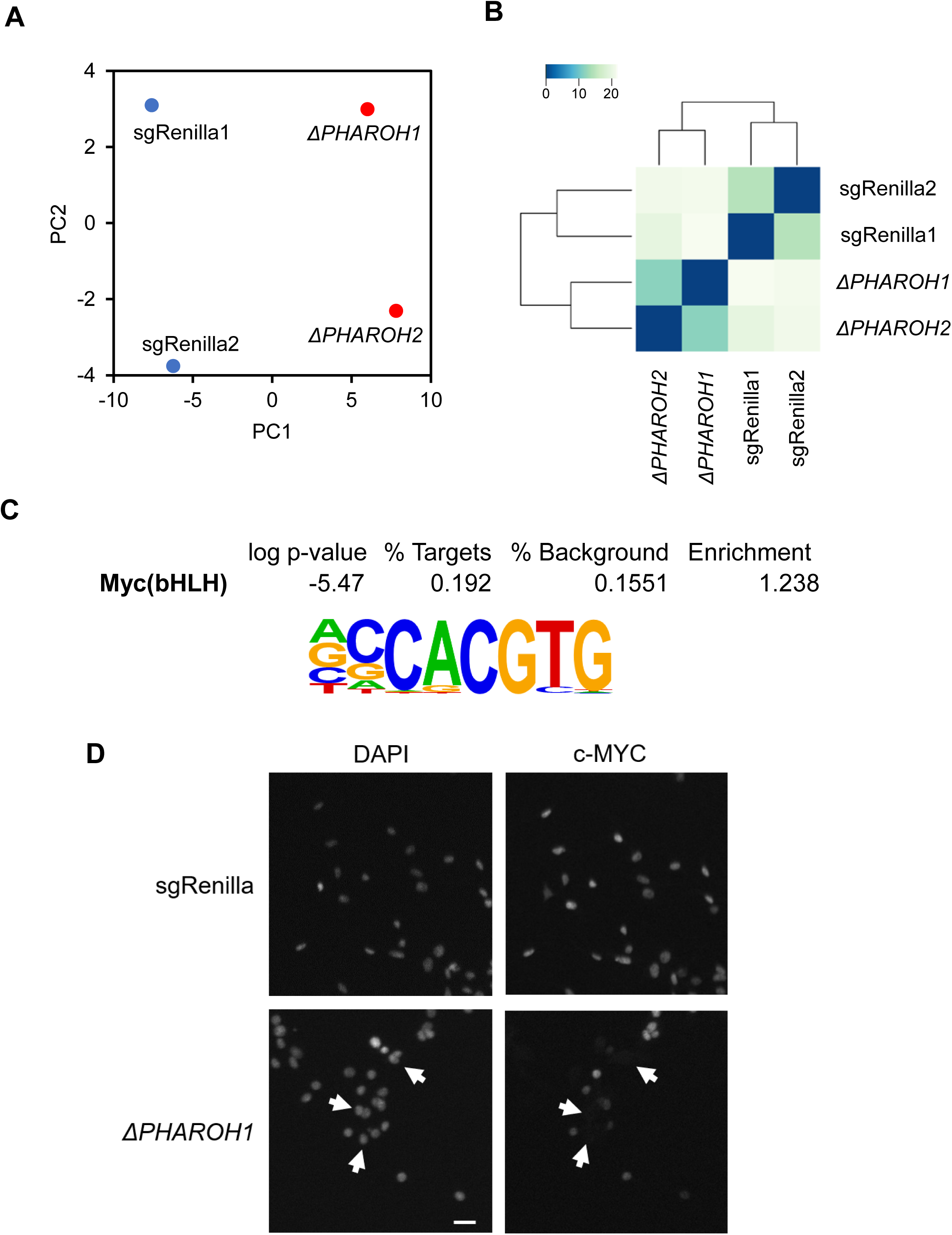
A. Principal component analysis of two sgRenilla negative control clones and two *PHAROH* knockout clones. Deletion of *PHAROH* is well separated by PC1. B. Euclidean distance plot indicating that the negative control clones and *PHAROH* knockout clones cluster independently. C. Motif analysis of promoter region of differentially expressed genes. c-MYC motif is enriched 1.24 fold over background sequences. D. Immunofluorescence of c-MYC in *PHAROH* knockout clones shows absence of c-MYC signal in a majority of cells. Scale bar = 50 μm

**Supplemental Figure 5.**
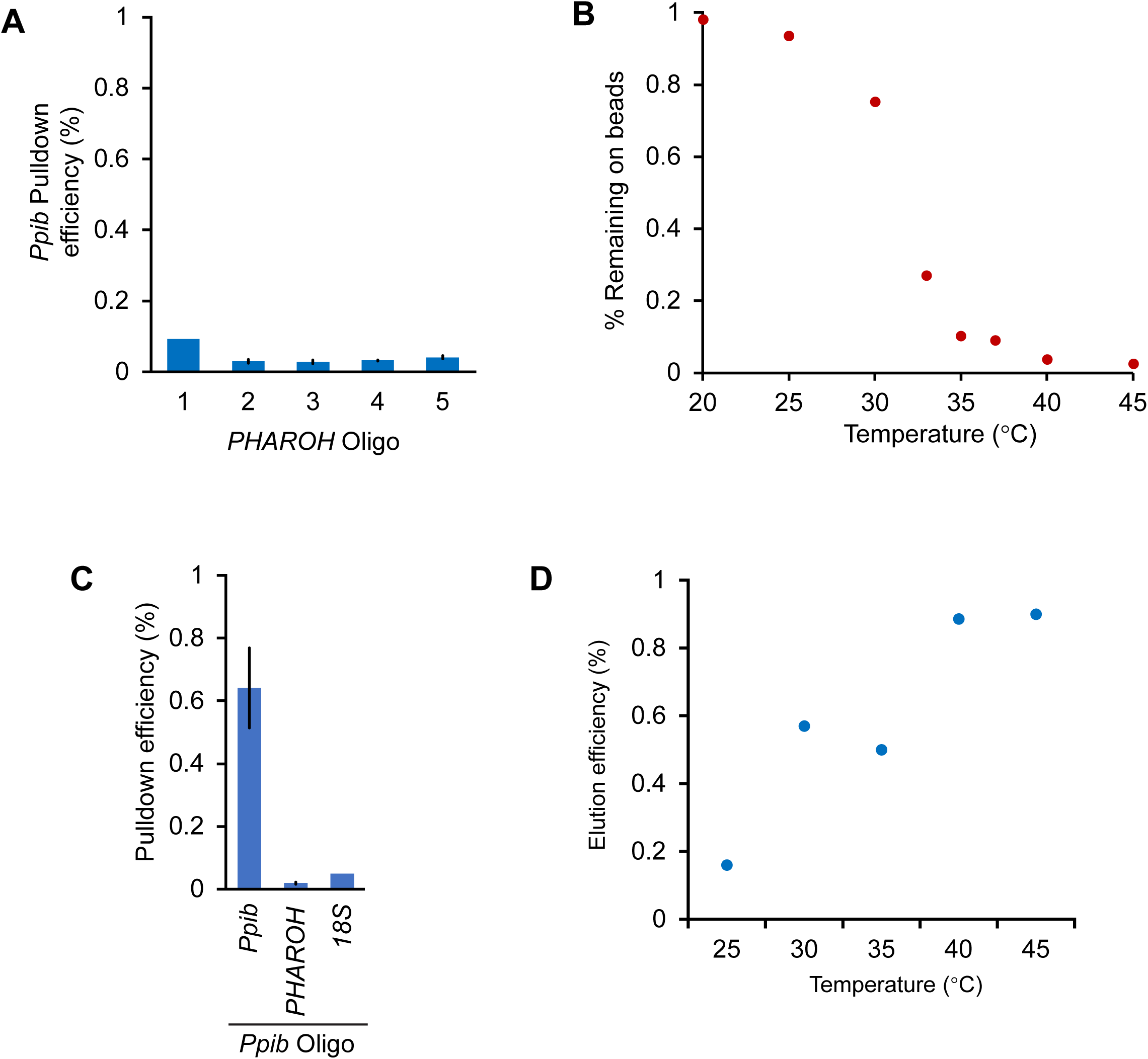
A. The amount of *PHAROH* RNA remaining on the beads after thermal elution is inverse to that of the eluate. B. Off-target pulldown of *Ppib* using *PHAROH* oligos is low. C. An oligo designed against *Ppib* can pull the RNA down at ∼65% efficiency, and does not pull down *PHAROH* or 18S. D. *Ppib* can also be eluted via a temperature gradient, and is optimally released at 40° C.

**Supplemental Figure 6.**
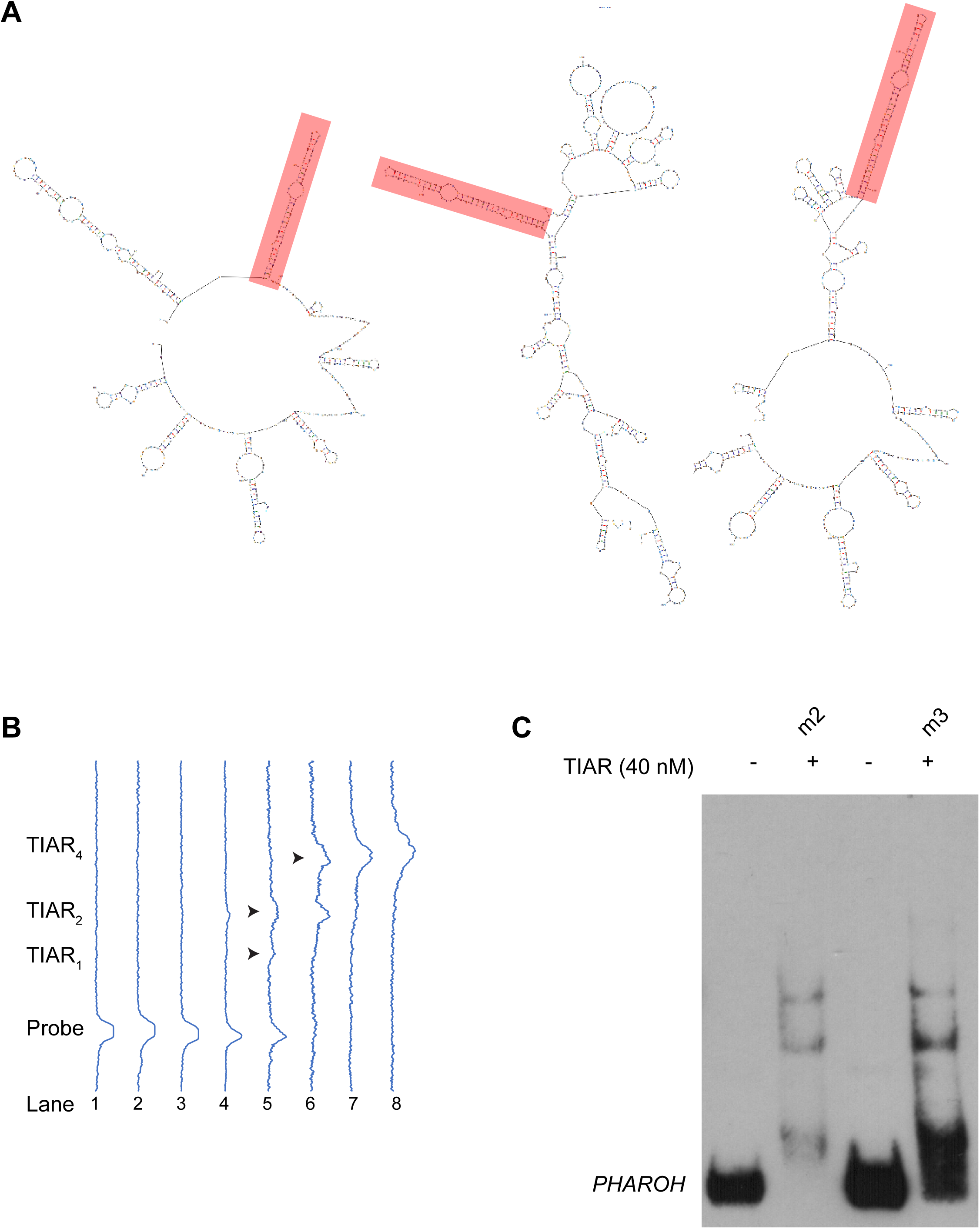
A. Mapping the top seven binding sites to predicted structures (top three shown here), reveals a conserved hairpin on the majority of predicted structures. B. Profile analysis of the RNA EMSA gel in Fig. 6C, showing the shift in intensity. C. Binding of TIAR to m2 and m3 are similar, possibly due to the mutation of a weaker binding site does not greatly impact overall binding.

**Supplemental Figure 7.**
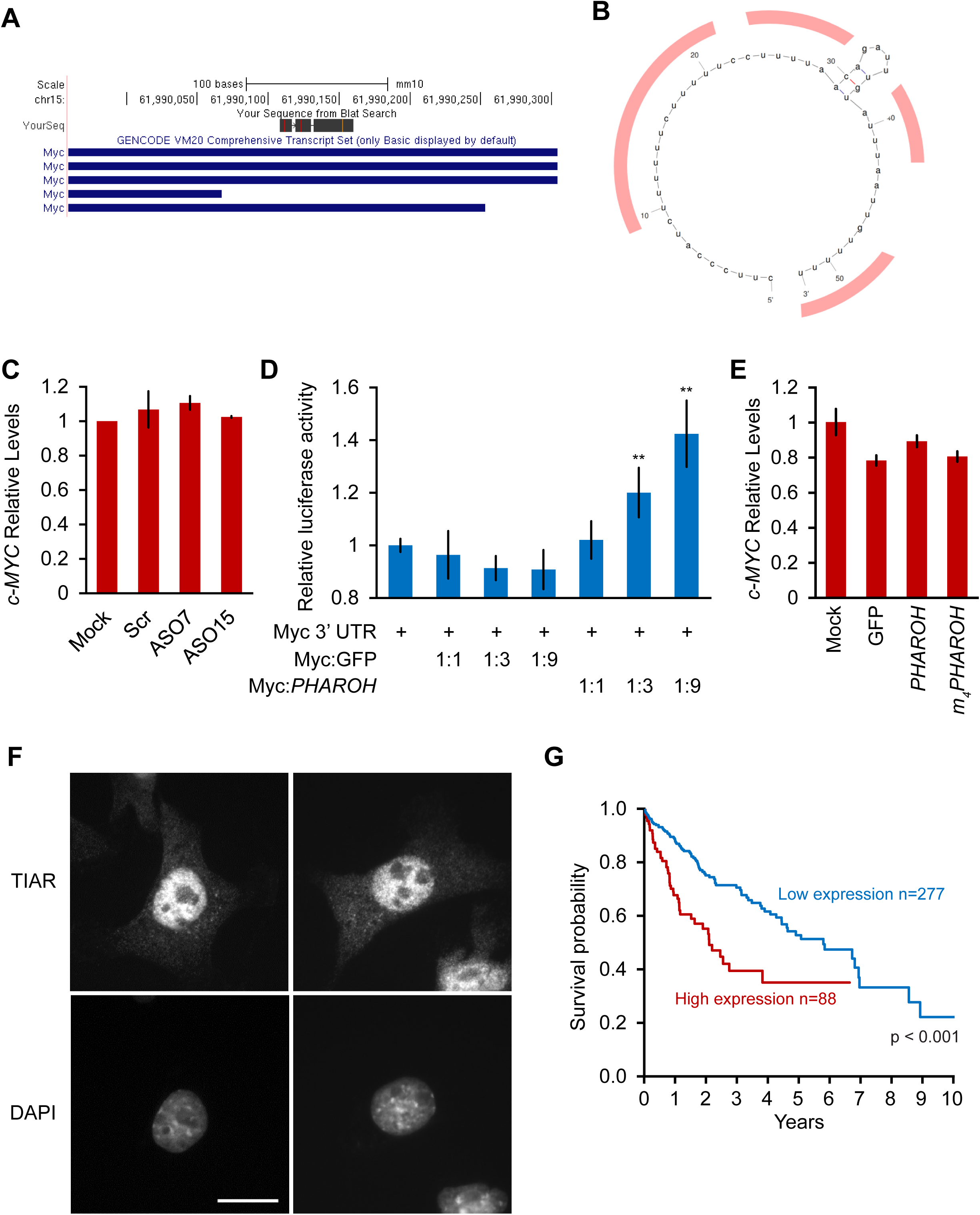
A. Of the two TIAR binding sites on c-MYC’s 3’ UTR, only one maps to the mouse genome. B. Potential TIAR binding sites on the mouse c-Myc 3’ UTR highlighted in red. C. Knockdown of *PHAROH* does not change c-Myc mRNA levels, suggesting that *PHAROH* acts at a post-transcriptional level. D. Addition of *PHAROH* to a luciferase construct with a c-Myc 3’ UTR increases luciferase activity in a dose dependent manner. E. c-MYC RNA levels do not change when *PHAROH* or TIAR are overexpressed. F. IF microscopy of TIAR showing predominantly nuclear localization. Scale bar = 25 μm G. Kaplan-Meier survival plot of patients with low and high TIAR expression.

## References

Allain, F. H., Bouvet, P., Dieckmann, T., & Feigon, J. (2000). Molecular basis of sequence-specific recognition of pre-ribosomal RNA by nucleolin. EMBO J, 19(24), 6870–6881. doi:10.1093/emboj/19.24.6870

Amodio, N., Raimondi, L., Juli, G., Stamato, M. A., Caracciolo, D., Tagliaferri, P., & Tassone, P. (2018). MALAT1: a druggable long non-coding RNA for targeted anti-cancer approaches. J Hematol Oncol, 11(1), 63. doi:10.1186/s13045-018-0606-4

Arun, G., Diermeier, S. D., & Spector, D. L. (2018). Therapeutic Targeting of Long Non-Coding RNAs in Cancer. Trends Mol Med, 24(3), 257–277. doi:10.1016/j.molmed.2018.01.001

Asrani, S. K., Devarbhavi, H., Eaton, J., & Kamath, P. S. (2019). Burden of liver diseases in the world. J Hepatol, 70(1), 151–171. doi:10.1016/j.jhep.2018.09.014

Berasain, C., Garcia-Trevijano, E. R., Castillo, J., Erroba, E., Lee, D. C., Prieto, J., & Avila, M. A. (2005). Amphiregulin: an early trigger of liver regeneration in mice. Gastroenterology, 128(2), 424–432. doi:10.1053/j.gastro.2004.11.006

Bergmann, J. H., Li, J., Eckersley-Maslin, M. A., Rigo, F., Freier, S. M., & Spector, D. L. (2015). Regulation of the ESC transcriptome by nuclear long noncoding RNAs. Genome Res, 25(9), 1336–1346. doi:10.1101/gr.189027.114

Cesana, M., Cacchiarelli, D., Legnini, I., Santini, T., Sthandier, O., Chinappi, M., … Bozzoni, I. (2011). A long noncoding RNA controls muscle differentiation by functioning as a competing endogenous RNA. Cell, 147(2), 358–369. doi:10.1016/j.cell.2011.09.028

Costa, F. F. (2005). Non-coding RNAs: new players in eukaryotic biology. Gene, 357(2), 83–94. doi:10.1016/j.gene.2005.06.019

Dhanasekaran, R., Nault, J. C., Roberts, L. R., & Zucman-Rossi, J. (2019). Genomic Medicine and Implications for Hepatocellular Carcinoma Prevention and Therapy. Gastroenterology, 156(2), 492–509. doi:10.1053/j.gastro.2018.11.001

Dinger, M. E., Amaral, P. P., Mercer, T. R., Pang, K. C., Bruce, S. J., Gardiner, B. B., … Mattick, J. S. (2008). Long noncoding RNAs in mouse embryonic stem cell pluripotency and differentiation. Genome Res, 18(9), 1433–1445. doi:10.1101/gr.078378.108

Djebali, S., Davis, C. A., Merkel, A., Dobin, A., Lassmann, T., Mortazavi, A., … Gingeras, T. R. (2012). Landscape of transcription in human cells. Nature, 489(7414), 101–108. doi:10.1038/nature11233

Dobin, A., Davis, C. A., Schlesinger, F., Drenkow, J., Zaleski, C., Jha, S., … Gingeras, T. R. (2013). STAR: ultrafast universal RNA-seq aligner. Bioinformatics, 29(1), 15–21. doi:10.1093/bioinformatics/bts635

Eng, J. K., McCormack, A. L., & Yates, J. R. (1994). An approach to correlate tandem mass spectral data of peptides with amino acid sequences in a protein database. J Am Soc Mass Spectrom, 5(11), 976–989. doi:10.1016/1044-0305(94)80016-2

Ezponda, T., & Licht, J. D. (2014). Molecular pathways: deregulation of histone h3 lysine 27 methylation in cancer-different paths, same destination. Clin Cancer Res, 20(19), 5001–5008. doi:10.1158/1078-0432.CCR-13-2499

Ferlay, J., Shin, H. R., Bray, F., Forman, D., Mathers, C., & Parkin, D. M. (2010). Estimates of worldwide burden of cancer in 2008: GLOBOCAN 2008. Int J Cancer, 127(12), 2893–2917. doi:10.1002/ijc.25516

Garcia-Irigoyen, O., Latasa, M. U., Carotti, S., Uriarte, I., Elizalde, M., Urtasun, R., … Avila, M. A. (2015). Matrix metalloproteinase 10 contributes to hepatocarcinogenesis in a novel crosstalk with the stromal derived factor 1/C-X-C chemokine receptor 4 axis. Hepatology, 62(1), 166–178. doi:10.1002/hep.27798

Greten, T. F., Lai, C. W., Li, G., & Staveley-O’Carroll, K. F. (2019). Targeted and Immune-Based Therapies for Hepatocellular Carcinoma. Gastroenterology, 156(2), 510–524. doi:10.1053/j.gastro.2018.09.051

Heinz, S., Benner, C., Spann, N., Bertolino, E., Lin, Y. C., Laslo, P., … Glass, C. K. (2010). Simple combinations of lineage-determining transcription factors prime cis-regulatory elements required for macrophage and B cell identities. Mol Cell, 38(4), 576–589. doi:10.1016/j.molcel.2010.05.004

Hoshida, Y., Nijman, S. M., Kobayashi, M., Chan, J. A., Brunet, J. P., Chiang, D. Y., … Golub, T. R. (2009). Integrative transcriptome analysis reveals common molecular subclasses of human hepatocellular carcinoma. Cancer Res, 69(18), 7385–7392. doi:10.1158/0008-5472.CAN-09-1089

Ji, J., Dai, X., Yeung, S. J., & He, X. (2019). The role of long non-coding RNA GAS5 in cancers. Cancer Manag Res, 11, 2729–2737. doi:10.2147/CMAR.S189052

Kedersha, N. L., Gupta, M., Li, W., Miller, I., & Anderson, P. (1999). RNA-binding proteins TIA-1 and TIAR link the phosphorylation of eIF-2 alpha to the assembly of mammalian stress granules. J Cell Biol, 147(7), 1431–1442. doi:10.1083/jcb.147.7.1431

Kim, H. S., Headey, S. J., Yoga, Y. M., Scanlon, M. J., Gorospe, M., Wilce, M. C., & Wilce, J. A. (2013). Distinct binding properties of TIAR RRMs and linker region. RNA Biol, 10(4), 579–589. doi:10.4161/rna.24341

Kim, H. S., Wilce, M. C., Yoga, Y. M., Pendini, N. R., Gunzburg, M. J., Cowieson, N. P., … Wilce, J. A. (2011). Different modes of interaction by TIAR and HuR with target RNA and DNA. Nucleic Acids Res, 39(3), 1117–1130. doi:10.1093/nar/gkq837

Lai, M. C., Yang, Z., Zhou, L., Zhu, Q. Q., Xie, H. Y., Zhang, F., … Zheng, S. S. (2012). Long non-coding RNA MALAT-1 overexpression predicts tumor recurrence of hepatocellular carcinoma after liver transplantation. Med Oncol, 29(3), 1810–1816. doi:10.1007/s12032-011-0004-z

Lee, S., Kopp, F., Chang, T. C., Sataluri, A., Chen, B., Sivakumar, S., … Mendell, J. T. (2016). Noncoding RNA NORAD Regulates Genomic Stability by Sequestering PUMILIO Proteins. Cell, 164(1-2), 69–80. doi:10.1016/j.cell.2015.12.017

Li, C., Chen, J., Zhang, K., Feng, B., Wang, R., & Chen, L. (2015). Progress and Prospects of Long Noncoding RNAs (lncRNAs) in Hepatocellular Carcinoma. Cell Physiol Biochem, 36(2), 423–434. doi:10.1159/000430109

Liao, B., Hu, Y., & Brewer, G. (2007). Competitive binding of AUF1 and TIAR to MYC mRNA controls its translation. Nat Struct Mol Biol, 14(6), 511–518. doi:10.1038/nsmb1249

Lin, R., Maeda, S., Liu, C., Karin, M., & Edgington, T. S. (2007). A large noncoding RNA is a marker for murine hepatocellular carcinomas and a spectrum of human carcinomas. Oncogene, 26(6), 851–858. doi:10.1038/sj.onc.1209846

Liu, L., Yue, H., Liu, Q., Yuan, J., Li, J., Wei, G., … Chen, R. (2016). LncRNA MT1JP functions as a tumor suppressor by interacting with TIAR to modulate the p53 pathway. Oncotarget, 7(13), 15787–15800. doi:10.18632/oncotarget.7487

Llovet, J. M., Montal, R., Sia, D., & Finn, R. S. (2018). Molecular therapies and precision medicine for hepatocellular carcinoma. Nat Rev Clin Oncol, 15(10), 599–616. doi:10.1038/s41571-018-0073-4

Love, M. I., Huber, W., & Anders, S. (2014). Moderated estimation of fold change and dispersion for RNA-seq data with DESeq2. Genome Biol, 15(12), 550. doi:10.1186/s13059-014-0550-8

Ma, W., & Mayr, C. (2018). A Membraneless Organelle Associated with the Endoplasmic Reticulum Enables 3’UTR-Mediated Protein-Protein Interactions. Cell, 175(6), 1492–1506 e1419. doi:10.1016/j.cell.2018.10.007

Mazan-Mamczarz, K., Lal, A., Martindale, J. L., Kawai, T., & Gorospe, M. (2006). Translational repression by RNA-binding protein TIAR. Mol Cell Biol, 26(7), 2716–2727. doi:10.1128/MCB.26.7.2716-2727.2006

McHugh, C. A., Chen, C. K., Chow, A., Surka, C. F., Tran, C., McDonel, P., … Guttman, M. (2015). The Xist lncRNA interacts directly with SHARP to silence transcription through HDAC3. Nature, 521(7551), 232–236. doi:10.1038/nature14443

Meyer, C., Garzia, A., Mazzola, M., Gerstberger, S., Molina, H., & Tuschl, T. (2018). The TIA1 RNA-Binding Protein Family Regulates EIF2AK2-Mediated Stress Response and Cell Cycle Progression. Mol Cell, 69(4), 622–635 e626. doi:10.1016/j.molcel.2018.01.011

Peng, S. Y., Lai, P. L., & Hsu, H. C. (1993). Amplification of the c-myc gene in human hepatocellular carcinoma: biologic significance. J Formos Med Assoc, 92(10), 866–870.

Perkins, D. N., Pappin, D. J., Creasy, D. M., & Cottrell, J. S. (1999). Probability-based protein identification by searching sequence databases using mass spectrometry data. Electrophoresis, 20(18), 3551–3567. doi:10.1002/(SICI)1522-2683(19991201)20:18<3551::AID-ELPS3551>3.0.CO;2-2

Philip, P. A., Mahoney, M. R., Allmer, C., Thomas, J., Pitot, H. C., Kim, G., … Erlichman, C. (2005). Phase II study of Erlotinib (OSI-774) in patients with advanced hepatocellular cancer. J Clin Oncol, 23(27), 6657–6663. doi:10.1200/JCO.2005.14.696

Piecyk, M., Wax, S., Beck, A. R., Kedersha, N., Gupta, M., Maritim, B., … Anderson, P. (2000). TIA-1 is a translational silencer that selectively regulates the expression of TNF-alpha. EMBO J, 19(15), 4154–4163. doi:10.1093/emboj/19.15.4154

Quintela-Fandino, M., Le Tourneau, C., Duran, I., Chen, E. X., Wang, L., Tsao, M., … Siu, L. L. (2010). Phase I combination of sorafenib and erlotinib therapy in solid tumors: safety, pharmacokinetic, and pharmacodynamic evaluation from an expansion cohort. Mol Cancer Ther, 9(3), 751–760. doi:10.1158/1535-7163.MCT-09-0868

Rimassa, L., & Santoro, A. (2009). Sorafenib therapy in advanced hepatocellular carcinoma: the SHARP trial. Expert Rev Anticancer Ther, 9(6), 739–745. doi:10.1586/era.09.41

Rinn, J. L., & Chang, H. Y. (2012). Genome regulation by long noncoding RNAs. Annu Rev Biochem, 81, 145–166. doi:10.1146/annurev-biochem-051410-092902

Ross, P. L., Huang, Y. N., Marchese, J. N., Williamson, B., Parker, K., Hattan, S., … Pappin, D. J. (2004). Multiplexed protein quantitation in Saccharomyces cerevisiae using amine-reactive isobaric tagging reagents. Mol Cell Proteomics, 3(12), 1154–1169. doi:10.1074/mcp.M400129-MCP200

Shu, L., Yan, W., & Chen, X. (2006). RNPC1, an RNA-binding protein and a target of the p53 family, is required for maintaining the stability of the basal and stress-induced p21 transcript. Genes Dev, 20(21), 2961–2972. doi:10.1101/gad.1463306

Siegel, R., Ma, J., Zou, Z., & Jemal, A. (2014). Cancer statistics, 2014. CA Cancer J Clin, 64(1), 9–29. doi:10.3322/caac.21208

Slaymaker, I. M., Gao, L., Zetsche, B., Scott, D. A., Yan, W. X., & Zhang, F. (2016). Rationally engineered Cas9 nucleases with improved specificity. Science, 351(6268), 84–88. doi:10.1126/science.aad5227

Soudyab, M., Iranpour, M., & Ghafouri-Fard, S. (2016). The Role of Long Non-Coding RNAs in Breast Cancer. Arch Iran Med, 19(7), 508–517. doi:0161907/AIM.0011

Tabula Muris, C., Overall, c., Logistical, c., Organ, c., processing, Library, p., … Principal, i. (2018). Single-cell transcriptomics of 20 mouse organs creates a Tabula Muris. Nature, 562(7727), 367–372. doi:10.1038/s41586-018-0590-4

Tani, H., Mizutani, R., Salam, K. A., Tano, K., Ijiri, K., Wakamatsu, A., … Akimitsu, N. (2012). Genome-wide determination of RNA stability reveals hundreds of short-lived noncoding transcripts in mammals. Genome Res, 22(5), 947–956. doi:10.1101/gr.130559.111

Tsai, M. C., Manor, O., Wan, Y., Mosammaparast, N., Wang, J. K., Lan, F., … Chang, H. Y. (2010). Long noncoding RNA as modular scaffold of histone modification complexes. Science, 329(5992), 689–693. doi:10.1126/science.1192002

Uhlen, M., Zhang, C., Lee, S., Sjostedt, E., Fagerberg, L., Bidkhori, G., … Ponten, F. (2017). A pathology atlas of the human cancer transcriptome. Science, 357(6352). doi:10.1126/science.aan2507

Villanueva, A. (2019). Hepatocellular Carcinoma. N Engl J Med, 380(15), 1450–1462. doi:10.1056/NEJMra1713263

Wang, J., Liu, X., Wu, H., Ni, P., Gu, Z., Qiao, Y., … Fan, Q. (2010). CREB up-regulates long non-coding RNA, HULC expression through interaction with microRNA-372 in liver cancer. Nucleic Acids Res, 38(16), 5366–5383. doi:10.1093/nar/gkq285

Yamada, N., Hasegawa, Y., Yue, M., Hamada, T., Nakagawa, S., & Ogawa, Y. (2015). Xist Exon 7 Contributes to the Stable Localization of Xist RNA on the Inactive X-Chromosome. PLoS Genet, 11(8), e1005430. doi:10.1371/journal.pgen.1005430

Yu, F. J., Zheng, J. J., Dong, P. H., & Fan, X. M. (2015). Long non-coding RNAs and hepatocellular carcinoma. Mol Clin Oncol, 3(1), 13–17. doi:10.3892/mco.2014.429

Yuan, S. X., Wang, J., Yang, F., Tao, Q. F., Zhang, J., Wang, L. L., … Zhou, W. P. (2016). Long noncoding RNA DANCR increases stemness features of hepatocellular carcinoma by derepression of CTNNB1. Hepatology, 63(2), 499–511. doi:10.1002/hep.27893

Zhao, R., Nakamura, T., Fu, Y., Lazar, Z., & Spector, D. L. (2011). Gene bookmarking accelerates the kinetics of post-mitotic transcriptional re-activation. Nat Cell Biol, 13(11), 1295–1304. doi:10.1038/ncb2341

Zheng, K., Cubero, F. J., & Nevzorova, Y. A. (2017). c-MYC-Making Liver Sick: Role of c-MYC in Hepatic Cell Function, Homeostasis and Disease. Genes (Basel), 8(4). doi:10.3390/genes8040123

